# Ceramide-induced Endoplasmic Reticulum Stress as a Targetable Vulnerability in Endocrine Therapy-Resistant Breast Cancer

**DOI:** 10.1101/2025.08.18.670862

**Authors:** Purab Pal, Shweta Chitkara, Godwin K. Sarpey, Fatimah Alani, Huiping Zhao, Malik Ata, Jun Qu, Rachel Schiff, Debra Tonetti, Geoffrey L. Greene, Jonna Frasor, Gunes Ekin Atilla-Gokcumen, Jonathan L. Coloff

## Abstract

Despite the success of endocrine therapy (ET) in treating hormone receptor-positive breast cancer, a significant proportion of patients relapse during or after treatment, making ET resistance a major clinical challenge. Previously we have shown that ET-resistant breast cancer cells exhibit reduced ceramide levels and an increased sensitivity to ceramide-induced cell death. Here, we demonstrate that ceramides induce a distinct transcriptional reprogramming in ET-resistant cells, characterized by upregulation of endoplasmic reticulum stress (EnRS) pathways. Ceramide-induced EnRS is PERK-dependent and functionally linked to cell death in multiple models of ET resistance. Using a photoactivatable ceramide probe, we identify TRAM1 as a functionally important ceramide-interacting protein (CIP) in ET-resistant cells that correlates with worse relapse-free survival and a more aggressive breast cancer phenotype in luminal breast cancer patients. Additionally, knockdown of TRAM1 phenocopies ceramide action in ET resistance, thereby suggesting its role in mediating ceramide-induced lethal actions in ET resistance. Together, our findings reveal that ET-resistant breast cancer cells are more sensitive to PERK-mediated EnRS as compared to their ET-sensitive counterparts. Ceramides can exploit this dependence by interacting with CIPs such as TRAM1, leading to PERK activation and consequential cell death preferentially in the ET-resistant breast cancer models.

## INTRODUCTION

Breast cancer accounts for approximately 30% of all cancer cases in women in the United States, making it one of the most common malignancies among middle-aged and older women.^1^ Around 80% of breast tumors express estrogen receptor-α (ER+).^2^ While hormone receptor-negative subtypes tend to be more aggressive and lethal,^3^ ER+ breast cancer remains responsible for the majority of breast cancer-related deaths.^3–5^

The current standard-of-care for ER+ breast cancer is endocrine therapy (ET) with or without CDK4/6 inhibitors. ET includes aromatase inhibitors or anti-estrogens such as Tamoxifen and Fulvestrant.^3, 6^ While most patients initially respond to ET, approximately 40% experience relapse either during treatment or after completing the standard five to ten-year adjuvant regimen. This suggests that ET resistance can arise either *de novo* or as a result of prolonged therapy.^7, 8^ Notably, most recurrent tumors retain ER expression but no longer respond to ET,^9^ highlighting the urgent need for improved therapeutic strategies for recurrent ER+ disease that are not based on the estrogen receptor.

ET-resistant cells undergo transcriptional^10–12^ and metabolic^13, 14^ reprogramming as a survival adaptation, which contributes to diminished therapeutic efficacy. Several studies have identified key metabolic alterations in ET resistance,^15^ including significant changes in cellular lipid composition.^16–18^ Our previous work demonstrated that ET-resistant breast cancer cells display an altered sphingolipid profile, marked by reduced ceramide levels compared to their ET-sensitive counterparts.^19^ Furthermore, ET-resistant cells show an increased sensitivity to ceramide-induced cell death, suggesting that ceramide depletion plays a functional role in ET-resistant cell survival.

Ceramides are well-established inducers of apoptosis, and their lethal effects have been extensively studied.^20, 21^ While ceramides can undergo a number of different interactions with mitochondrial lipids and proteins triggering the intrinsic apoptotic cascade,^21, 22^ recent studies have identified alternative mitochondria-independent mechanisms of ceramide-induced cell death,^23, 24^ such as endoplasmic reticulum stress (EnRS),^25–27^ autophagic cell death,^28, 29^ metabolic inhibition,^30^ and lipid peroxidation-mediated ferroptosis.^31, 32^ While lipid peroxidation-induced ferroptosis has been suggested as a potential vulnerability to target ET-resistant cells,^33^ we comprehensively analyzed the molecular mechanisms underscoring ceramide’s differential cytotoxicity in ET resistance.

Here, we describe our investigation of the molecular basis of ceramide’s increased lethality in ET-resistant breast cancer. By integrating lipidomic, transcriptomic, and proteomic results, we demonstrate that ceramides preferentially trigger lethal EnRS *via* activation of the PERK pathway in ET-resistant breast cancer models. We also found that ET-resistant breast cancer cells have an increased sensitivity to PERK-mediated EnRS compared to their parental ET-sensitive counterparts. Ceramides exploit this vulnerability by interacting with proteins such as TRAM1, which is critical for the survival of ET-resistant cells. Through these interactions, ceramides induce lethal levels of EnRS in ET-resistant cells. Therefore, our findings reveal a novel vulnerability in ET-resistant cells offering potential opportunities for therapeutic intervention specifically targeting ET-resistant BCs.

## RESULTS

### Ceramides selectively affect the transcriptome of ET-resistant models

We previously reported that ET-resistant breast cancer cells exhibit reduced ceramide levels and an increased sensitivity to ceramide-induced cell death.^19^ Here, we show that several ceramide species are significantly downregulated in multiple ET-resistant cell line models^34–36^ (**Fig S1A, S1B**). ET-resistant cells also show greater sensitivity to exogenous ceramide addition (**Fig S1C**) and to endogenous ceramide accumulation induced by ceramide kinase (CERK) inhibition^19^ (**Fig S1D**), suggesting that the reduced ceramide levels in ET-resistant cells are functionally relevant.

Ceramides are important signaling molecules that are known to influence numerous cellular processes.^37–40^ To investigate the differential actions of C8-ceramide in ET resistance, we performed total RNA-seq on ET-sensitive MCF-7 cells and three of its ET-resistant derivatives (MCF-7-HER2, MCF-7-TAMR, and MCF-7-FULR)^34, 35^ treated with or without C8-ceramide, the delivery of which we confirmed by LC-MS/MS (**Fig S2**). Principal component analysis (PCA) reveals that all ET-resistant lines differ substantially from parental MCF-7 cells, and that ceramide induces greater transcriptomic changes in ET-resistant models, whereas its effect on parental MCF-7 cells is minimal (**Fig 1A**). Specifically, C8-ceramide altered 252 genes in MCF-7, while it altered 4326, 2552, and 862 genes in MCF-7-HER2, MCF-7-TAMR, and MCF-7-FULR, respectively (Fold Change>2, P<0.05, FDR<0.01) (**Fig 1B** and **Table S1**).

**Figure 1:**
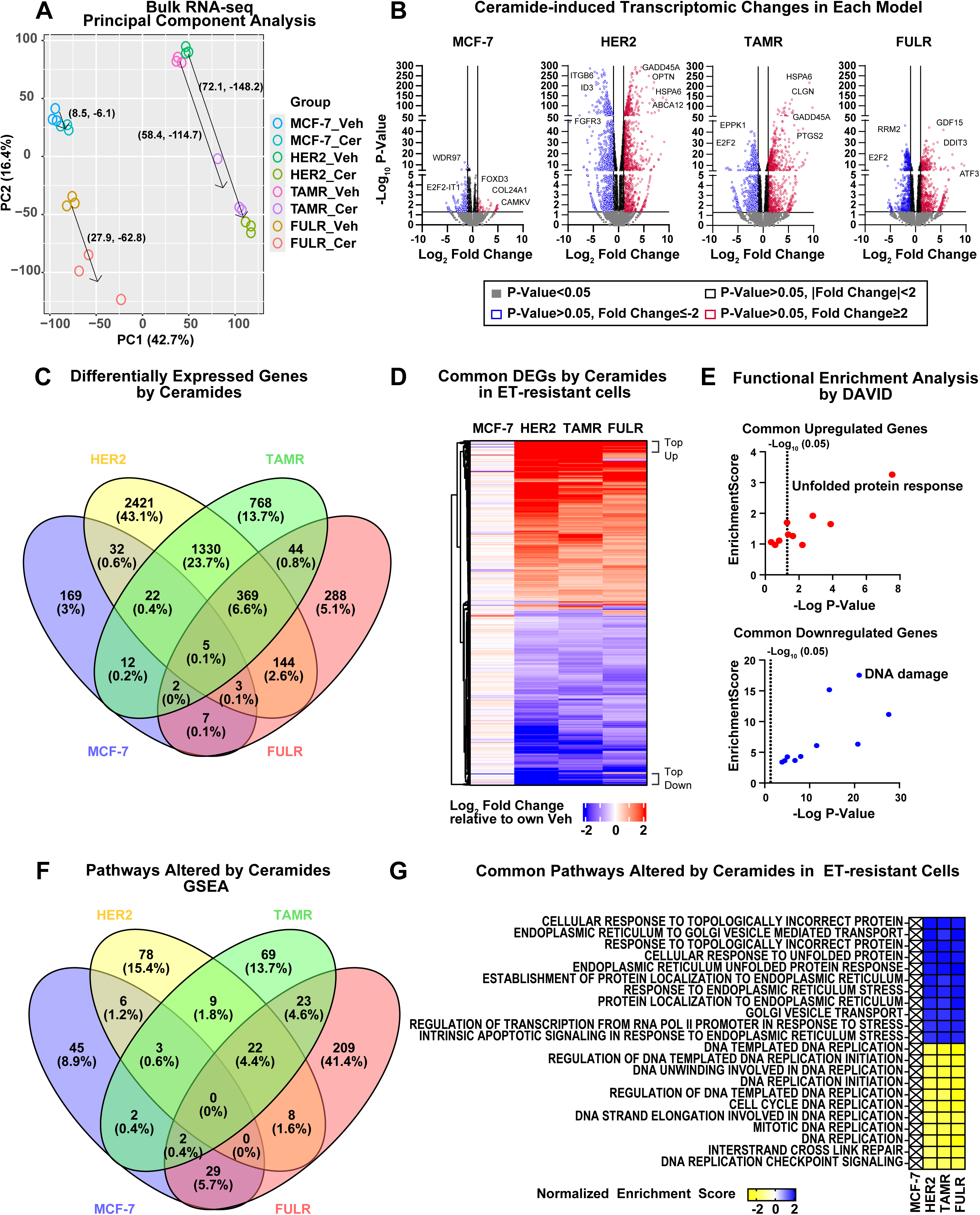
C8-ceramide preferentially causes endoplasmic reticulum stress in ET-resistant models. ET-sensitive MCF-7 cells and three ET-resistant derivatives were treated with 10 µM C8-ceramide (Cer) or ethanol as vehicle for 24 hours, followed by RNA-seq. **A.** Principal Component Analysis (PCA) plot of gene expression data obtained *via* RNA-seq (n=3 biological replicates). Arrows indicate ceramide-induced distances in each model. **B.** Volcano plots showing differentially expressed genes (DEGs) following C8-ceramide treatment in each model (fold change >2, P <0.05, FDR <0.01). **C.** Number of DEGs following C8-ceramide treatment in each model, shown as a Venn diagram. **D.** Hierarchical clustering of common DEGs across all ET-resistant models represented as a heatmap of mean Log_2_ fold change relative to vehicle-treated controls. **E.** Functional enrichment analysis of the top upregulated and downregulated genes (*via* DAVID); top 10 clusters (ranked by enrichment score) are shown. **F.** Venn diagram of Gene Set Enrichment Analysis (GSEA) comparing ceramide-vs. vehicle-treated samples in each model. **G.** The 22 pathways commonly altered across all ET-resistant models are displayed as a heatmap of normalized enrichment scores.

To identify pathways differentially affected by ceramides in ET-resistant cells, we first analyzed all differentially expressed genes (DEGs), where we found 369 genes that are commonly altered by C8-ceramide across all ET-resistant models but not in parental MCF-7 cells (**Fig 1C**). Hierarchical clustering of these DEGs identified top up- and down-regulated genes (**Fig 1D**), which we validated by RT-qPCR (**Fig S3**). Functional enrichment analysis (DAVID) showed that upregulated DEGs are significantly associated with the unfolded protein response (UPR) and EnRS, while the downregulated DEGs are linked to DNA damage and repair pathways (**Fig 1E** and **Table S2**). We also performed Gene Set Enrichment Analysis (GSEA) individually for each cell line, treated with or without C8-ceramide (Nom P<0.05, FDR<0.1) (**Table S3**) where we found 22 commonly altered pathways that were not altered in parental MCF-7 cells (**Fig 1F**). This approach revealed similar results, with upregulated pathways linked to EnRS and downregulated pathways associated with DNA replication (**Fig 1G**). These findings suggest that C8-ceramide dramatically reprograms the ET-resistant transcriptome while having a minimal impact on ET-sensitive cells.

### Elevated ISR in ET resistance is associated with an increased sensitivity to PERK activation

We and others have previously demonstrated that prolonged ET activates the Integrated Stress Response (ISR) to promote survival in ER+ breast cancer cells.^41–46^ Indeed, we found that ISR is among the top altered pathways across all ET-resistant models (**Fig 2A, B** and **Table S4**). The ISR pathway integrates stress signals from multiple stressors such as amino acid starvation, viral transfection, EnRS, and heme depletion.^47^ Since EnRS is one of the top pathways induced by ceramide in all resistant models, we investigated if ET-activated ISR is dependent on EnRS.

**Figure 2:**
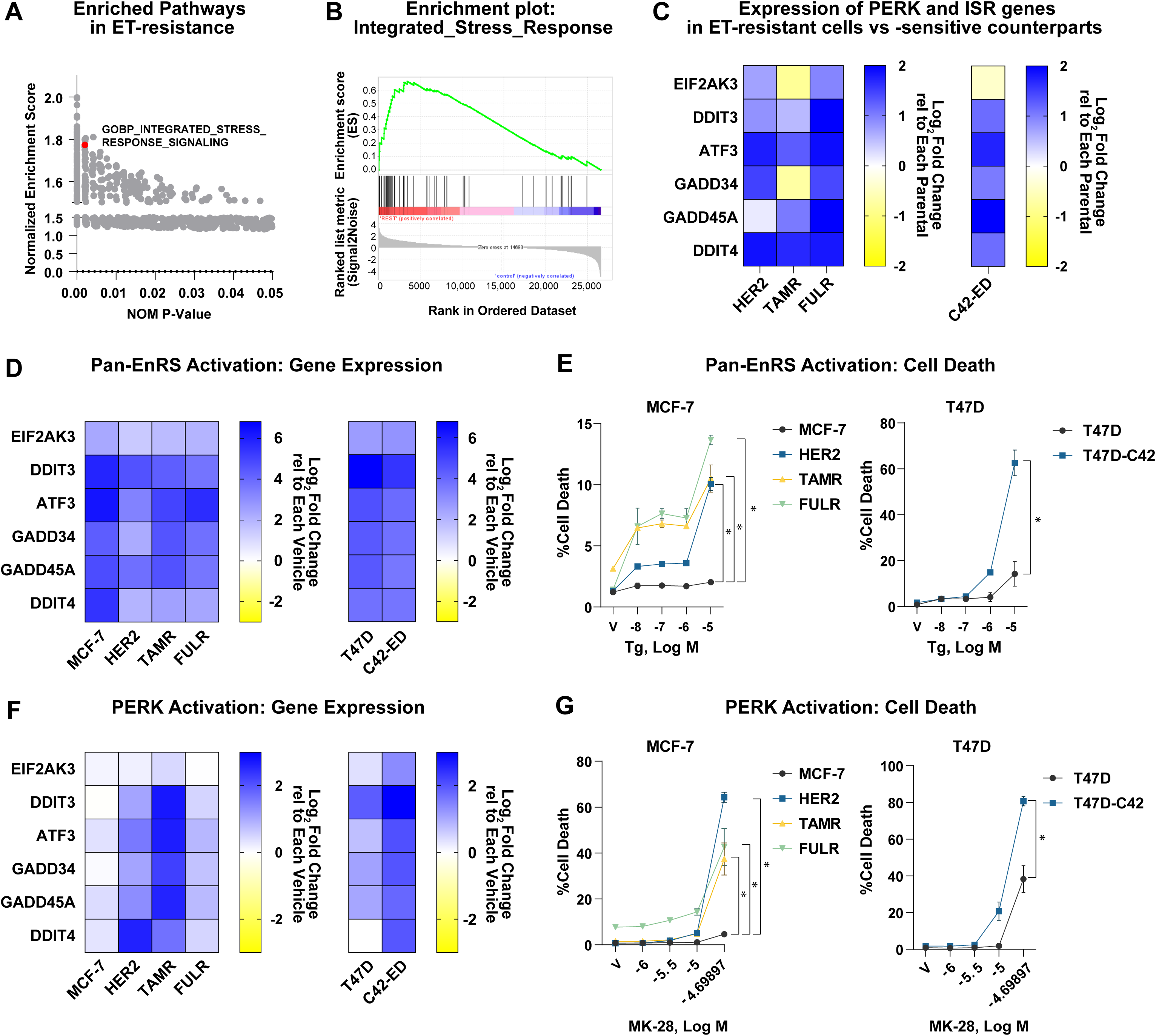
Elevated ISR is a vulnerability in multiple models of ET resistance. **A-B.** Top GSEA pathways comparing parental MCF-7 cells to three ET-resistant derivatives. **C.** RT-qPCR of ISR genes in ET-resistant derivatives of MCF-7 cells and in the long-term estrogen-deprived (LTED) ET-resistant derivative of parental T47D cells. Relative gene expression is shown as Log_2_ fold change (mean, n=3) relative to corresponding ET-sensitive parental cell lines. **D.** RT-qPCR of ISR genes from parental and ET-resistant MCF-7 and T47D cells treated with or without 1 µM Thapsigargin (Tg) for 24 hours. Data are represented as mean Log_2_ fold change with Thapsigargin relative to vehicle-treated controls in each cell line (mean, n=3). **E.** Cell death assessed by Hoechst-PI staining from cells treated with increasing concentrations of Thapsigargin (Tg) for 48 hours (V= vehicle-treated control). Values are the mean *±* SEM for triplicate samples from an experiment representative of two independent experiments. *P<0.05 by two-way ANOVA with Geisser-Greenhouse correction. The dose response of MCF-7 is compared to that of each resistant cell line. **F.** RT-qPCR from cells treated with 10 µM MK-28, a small-molecule activator of PERK, for 24 hours. Data are represented as mean Log_2_ fold change relative to vehicle-treated controls in each cell line (mean, n=3). **G.** Cell death assessed by Hoechst-PI staining from cells treated with increasing concentrations of MK-28 for 48 hours (V= vehicle-treated control). Values are the mean *±* SEM for triplicate samples from an experiment representative of two independent experiments. *P<0.05 by two-way ANOVA with Geisser-Greenhouse correction. The dose response of MCF-7 is compared to that of each resistant cell line.

Of the ISR-responsive genes,^47^ *EIF2AK3* (PERK, the ISR kinase that is activated by EnRS), *HSPA5* (BiP), *ATF4*, and *DDIT3* (CHOP) were present in the gene list that drives the ISR signature enrichment in ET resistance (**Table S5**). RT-qPCR data confirmed the elevated expression of these ISR genes in multiple ET-resistant cell lines as compared to their ET-sensitive counterparts (**Fig 2C** and **Fig S4A, B**).

To determine whether the pre-existing ISR signaling alters the sensitivity of ET-resistant cells to EnRS and PERK activation, we treated ET-sensitive and -resistant cells with Thapsigargin, a pan-EnRS activator and MK-28, a selective PERK activator.^48^ Thapsigargin similarly activates ISR genes in all cell lines (**Fig 2D** and **Fig S4C, D**) but preferentially induces cell death in ET-resistant models (**Fig 2E**), suggesting that ET-resistant cells are more sensitive to EnRS-induced cell death. In contrast, MK-28 induces a more pronounced ISR activation in ET-resistant cells as compared to ET-sensitive cells (**Fig 2F** and **Fig S4E, F**) and causes significantly higher cell death (**Fig 2G**), indicating that ET-resistant cells exhibit an elevated susceptibility to PERK activation and PERK-dependent cell death.

To test whether ISR sensitivity in ET-resistant cells is EnRS and PERK-specific, we induced ISR through amino acid starvation, which is known to activate another ISR kinase, *EIF2AK4* (GCN2). Indeed, we found that amino acid starvation induces ISR genes in all cell lines, similar to Thapsigargin (**Fig S5A**), but does not significantly alter cell death (**Fig S5B**) or suppress growth differently between ET-sensitive and ET-resistant models (**Fig S5C**). Collectively, these findings suggest that prolonged ET exposure primes a PERK-dependent ISR program that renders ET-resistant cells selectively vulnerable to further EnRS, PERK activation, and PERK-mediated cell death.

### Ceramide-induced PERK activation is lethal for ET-resistant models

We next examined the extent to which ceramide induces EnRS and PERK activation in ET-sensitive and ET-resistant cells. C8-ceramide addition increases ISR gene expression within 6 hours post-treatment in ET-resistant cells while having a minimal effect on ET-sensitive cells (**Fig 3A, B** and **Fig S6A, B**). Western blot analysis confirmed that C8-ceramide addition activates PERK (as indicated by its upward shift), stabilizes ATF4, and induces its pro-apoptotic target *DDIT3* (CHOP) specifically in ET-resistant models (**Fig 3C, D**).

**Figure 3:**
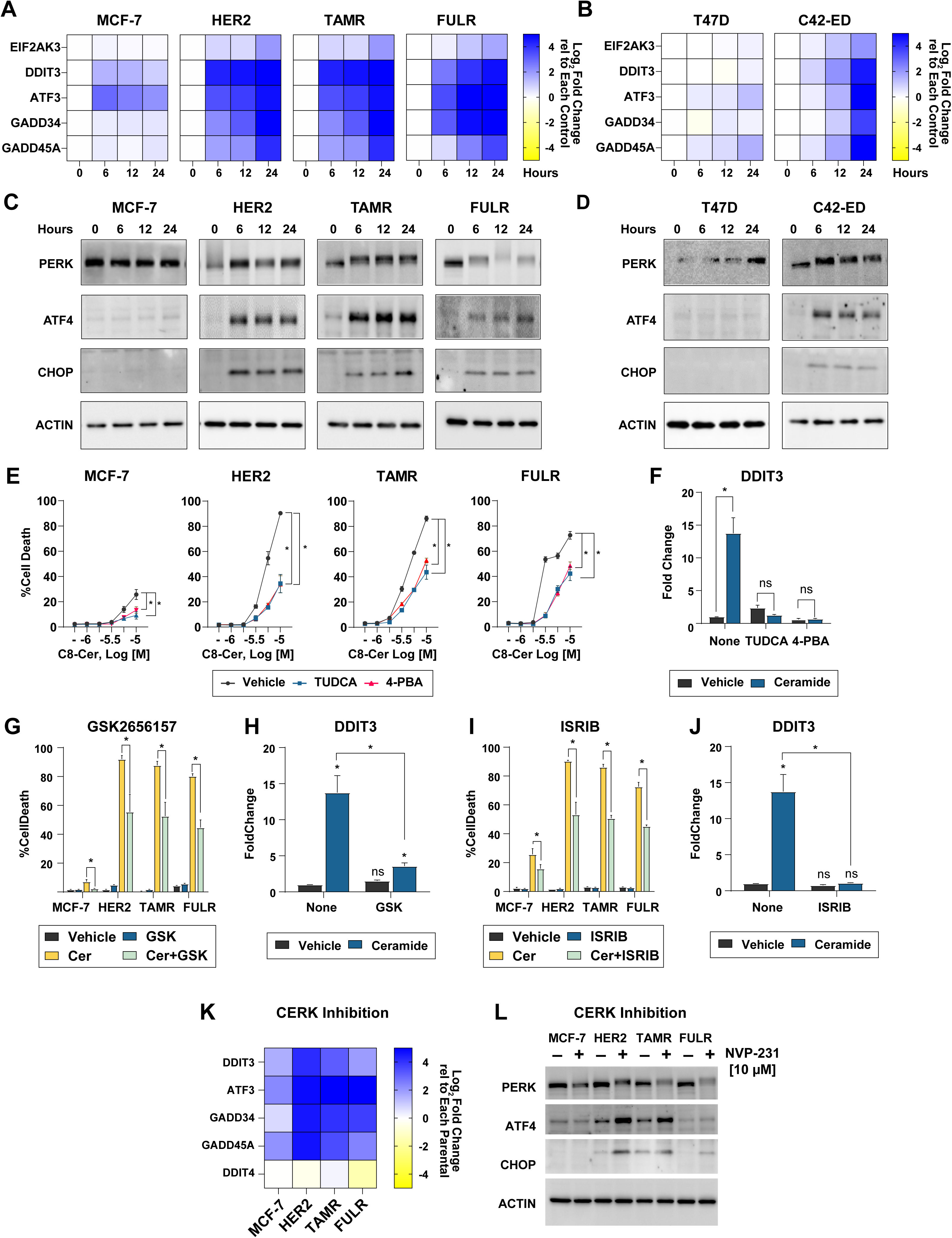
Ceramide induces lethal endoplasmic reticulum stress *via* PERK signaling. **A-D.** RT-qPCR of ISR genes from cells cultured with 10 µM C8-ceramide for up to 24 hours. Data are presented as mean Log_2_ fold change relative to vehicle-treated controls in each cell line (n=3) from one experiment representative of three independent experiments (**A-B**). Western blots showing protein expression levels of PERK, ATF4, and CHOP (representative of at least two independent experiments) (**C-D**). **E.** Cell death assessed by Hoechst-PI staining from cells pretreated with 100 µM Tauroursodeoxycholic acid (TUDCA) and 100 µM 4-phenylbutyric acid (4-PBA) for 2 hours, followed by C8-ceramide treatment for 46 hours. The values are the mean *±* SEM for triplicate samples from an experiment representative of two independent experiments. *P<0.05 by two-way ANOVA with Geisser-Greenhouse correction. **F.** RT-qPCR from MCF-7-TAMR cells pretreated with 100 µM TUDCA and 100 µM 4-PBA for 2 hours, followed by 5 µM C8-ceramide treatment for 22 hours. Data are presented as mean *±* SEM for triplicate samples from one experiment. *P<0.05 by unpaired Student’s t-test with Welch’s correction. **G.** Cell death assessed by Hoechst-PI staining from cells cultured with or without 10 µM C8-Ceramide, either alone or in combination with 10 µM GSK2656257 (GSK, PERK inhibitor), for 48 hours. Data are presented as mean *±* SEM for triplicate samples from an experiment representative of two independent experiments. *P<0.05 by unpaired Student’s t-test with Welch’s correction. **H.** RT-qPCR from MCF-7-TAMR cells treated with or without 5 µM C8-ceramide in combination with 10 µM GSK2656157 for 24 hours. Data are presented as mean *±* SEM for triplicate samples from one experiment. *P<0.05 by unpaired Student’s t-test with Welch’s correction. **I.** Cell death assessed by Hoechst-PI staining from cells pretreated with or without 20 µM ISRIB for 2 hours, followed by 46 hours of treatment with 10 µM C8-ceramide. Data are presented as mean *±* SEM for triplicate samples from an experiment representative of two independent experiments. *P<0.05 by unpaired Student’s t-test with Welch’s correction. **J.** RT-qPCR from MCF-7-TAMR cells pretreated with 20 µM ISRIB, followed by 22 hours of 5 µM C8-ceramide treatment. Data are presented as mean *±* SEM for triplicate samples from one experiment. *P<0.05 by unpaired Student’s t-test with Welch’s correction. **K.** RT-qPCR of ISR genes from cells cultured with 10 µM NVP-231 for 24 hours. Data are presented as mean Log_2_ fold change relative to vehicle-treated controls in each cell line (n=3) from one experiment representative of two independent experiments. **L.** Western blots showing PERK, ATF4, and CHOP expression following 24 hours of 10 µM NVP-231 treatment in MCF-7 parental and ET-resistant cells (representative of three independent experiments).

Since EnRS can be either pro-apoptotic or pro-survival,^49^ we used small molecules to assess the functional consequences of ceramide-induced EnRS in ET resistance. Inhibiting EnRS with Tauroursodeoxycholic acid (TUDCA) and 4-phenylbutyric acid (4-PBA) partially reverses ceramide-induced cell death (**Fig 3E**) alongside a decrease in *DDIT3* expression (**Fig 3F**). Similarly, inhibiting PERK with GSK2656157 or blocking downstream phospho-eIF2α signaling with ISRIB partially reverses ceramide-induced cell death (**Fig 3G, I**) and *DDIT3* gene expression (**Fig 3H, J**). Additionally, treatment with Salubrinal which prevents phospho-eIF2α dephosphorylation, enhances ceramide-induced cell death in ET-resistant cells (**Fig S7**).

As a complimentary approach to the direct addition of C8-ceramide, we also treated cells with NVP-231, an inhibitor of ceramide kinase (CERK), which we have previously shown to increase endogenous ceramide levels in both ET-sensitive and ET-resistant cells.^19^ Similar to C8-ceramide, we found that NVP-231 causes a stronger PERK activation in ET-resistant cells as compared to their parental ET-sensitive cells (**Fig 3K, L** and **Fig S6C**). Together, these findings demonstrate that ceramide-induced PERK-mediated EnRS selectively trigger cell death in ET-resistant cell line models.

### Ceramide-interacting proteins reveal potential upstream mediators of ceramide action in ET-resistance

To identify potential molecular mediators of ceramide’s unique action in ET resistant cells, we investigated ceramide-protein interactions in ET-resistant and -sensitive cells. Given the established role of ceramides as bioactive lipids that can directly interact with proteins,^50–52^ we hypothesized that distinct ceramide-protein interactions might underpin the unique responses to ceramides as observed in the ET-resistant contexts. To test this hypothesis, we utilized a photoactivatable and clickable C16:0 ceramide analog (pacCer) to capture ceramide-interacting proteins (CIPs)^53, 54^ from ET-sensitive parental MCF-7 and their ET-resistant derivative, MCF-7-HER2 cell lysates. Subsequent quantitative proteomic analyses enabled a comparative assessment of CIP abundance between the two cell lines (**Fig 4A**). We identified 948 CIPs significantly enriched and 274 CIPs downregulated in MCF-7-HER2 cells relative to MCF-7 parental cells (**Fig 4B** and **Table S6**). Importantly, the differential abundance of CIPs between the two lines closely mirrored their corresponding gene expression profiles (**Fig 4C**), supporting the identification of differentially expressed CIPs in the ET-resistant counterpart. Functional enrichment analyses revealed that the enriched CIPs are involved in activating cellular pathways such as EIF2 signaling, and Actin cytoskeleton signaling, whereas the downregulated cellular pathways consist of EnRS among various other pathways (**Fig 4D** and **Table S7**).

**Figure 4:**
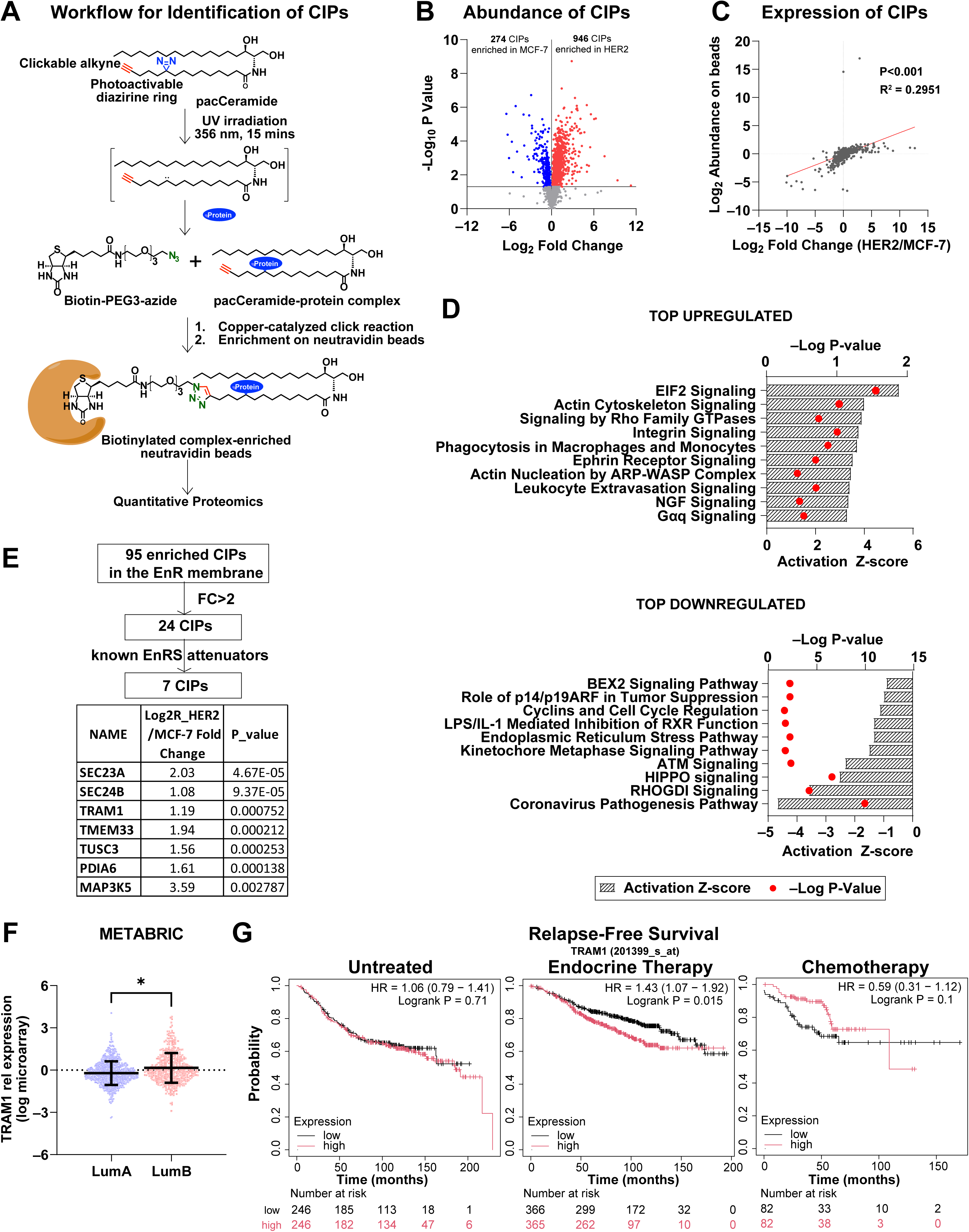
Profiling of ceramide-interacting proteins (CIPs) reveals TRAM1 as a potential mediator of ceramide action. **A.** Schematic diagram showing the workflow for identifying ceramide-interacting proteins (CIPs). **B.** Volcano plot showing differential enrichment of CIPs between parental MCF-7 and ET-resistant MCF-7-HER2 cells. **C.** Scatter plot of relative CIP abundance in MCF-7-HER2 vs. parental MCF-7 cells (Y-axis) plotted against mRNA expression levels of the same CIPs from RNAseq data (X-axis). **D.** Ingenuity Pathway Analysis (IPA) of enriched CIPs in MCF-7-HER2 cells, revealing the top upregulated and downregulated pathways. **E.** Enriched CIPs localized to the endoplasmic reticulum in MCF-7-HER2 cells were sorted by fold enrichment and known function to identify a panel associated with attenuation of EnRS. **F.** Percentage of ER+ breast cancer patients on ET with luminal A or luminal B subtype, stratified by TRAM1-high and TRAM1-low cohorts in the METABRIC dataset. **G.** Kaplan-Meier plots showing relapse-free survival of breast cancer patients when untreated or treated with ET or chemotherapy, stratified by TRAM1-high and TRAM1-low cohorts. Cohorts were defined by median TRAM1 mRNA expression level in the kmPlotter database.

Given the importance of EnRS in the response to ceramides, we focused on CIPs that might play a role in attenuating EnRS and promoting survival in ET-resistant cells. Therefore, we selected CIPs that are located in the EnR, exhibit greater than 2-fold enrichment in MCF-7-HER2 cells, and have previously been implicated in the attenuation of EnRS (**Fig 4E**). This resulted in seven proteins referred to as EnR-CIPs.^55–61^

Among the 7 EnR-CIPs that came out of this analysis, only TRAM1 gene expression demonstrated a consistent and significant association with poor relapse-free survival of patients across multiple datasets (**Fig S8**). High TRAM1 expression was also associated with a luminal B phenotype in ET-treated patients (**Fig 4F**) and predicted poorer relapse-free survival in the ER+ breast cancer patient cohort that received ET, but not those that received chemotherapy or were untreated (**Fig 4G**), suggesting that TRAM1 may play a role in ET resistance.

### TRAM1 mediates ceramide action in ET-resistant models

TRAM1 (Translocation Associated Membrane Protein 1) is a component of the EnR-translocon complex, that facilitates the import of nascent polypeptide chains into the EnR lumen and mediates the retrograde translocation of misfolded or unfolded proteins to the cytosol (**Fig 5A**).^57, 62, 63^ To investigate a potential role for TRAM1 in ET resistance, first we profiled TRAM1 expression in both ET-sensitive and ET-resistant MCF-7 cells, where we observed that TRAM1 expression is elevated across all ET-resistant derivatives of MCF-7 cells (**Fig 5B**). Silencing TRAM1 by siRNA (**Fig 5C**) results in increased cell death in all ET-resistant cells while having a minor impact on parental controls (**Fig 5D**). In addition, TRAM1 knockdown leads to a more pronounced upregulation of ISR genes in all ET-resistant models compared to the parental cells (**Fig 5E** and **Fig S9A**) suggesting TRAM1 inhibition phenocopies the preferential PERK activation by ceramides in ET-resistant cells. Notably, TRAM1 knockdown induces gene expression changes that correlate with top ceramide-induced DEGs in all the ET-resistant MCF-7 cells but not in the parental ET-sensitive MCF-7 cells (**Fig 5F** and **Fig S9B**), implicating TRAM1 to be a potential mediator of ceramide action in ET resistance.

**Figure 5:**
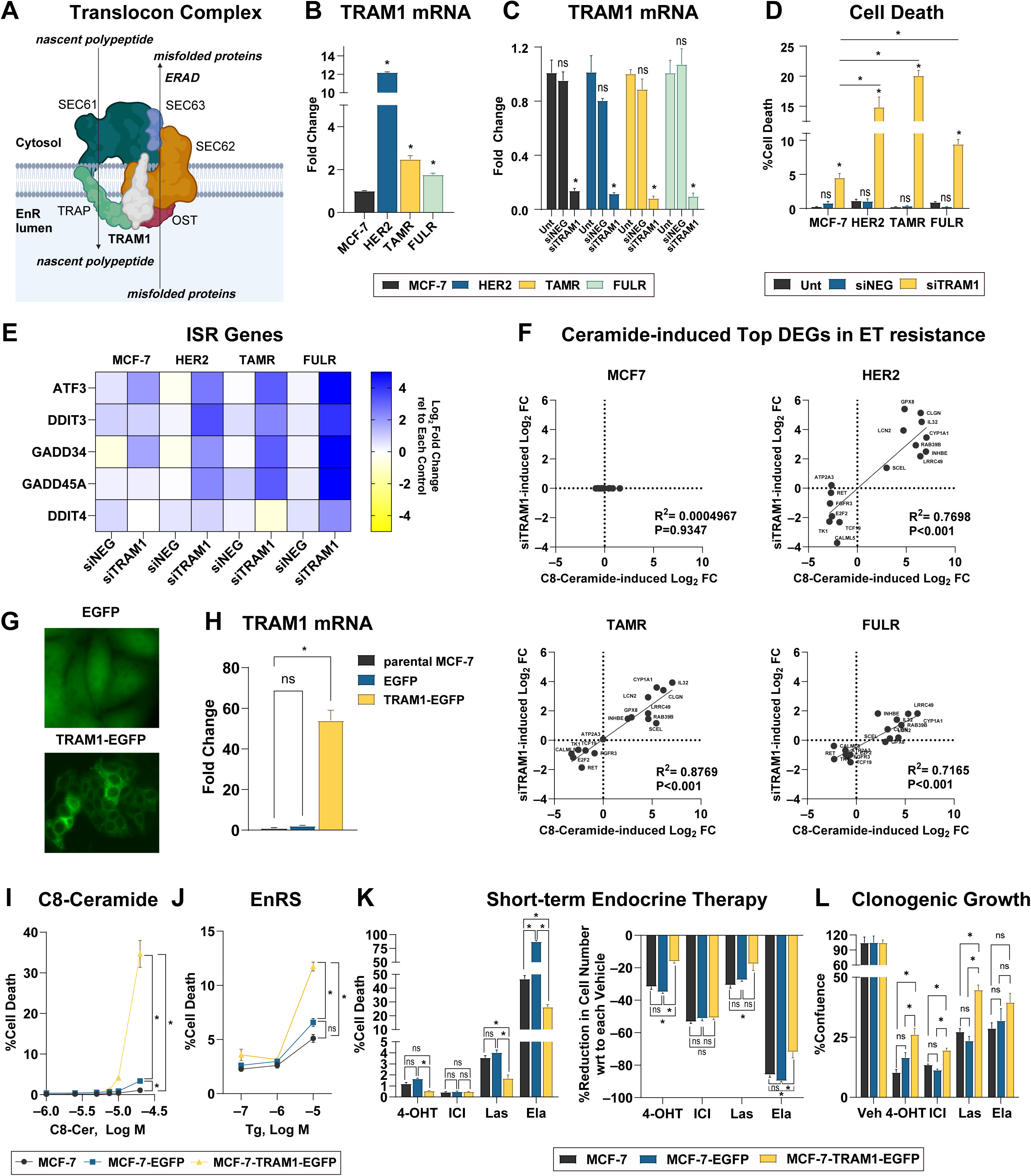
TRAM1 is a potential mediator of ceramide action in ET resistance. **A.** Schematic diagram of TRAM1 function. TRAM1 is a component of the endoplasmic reticulum (EnR) translocon complex. **B.** RT-qPCR of TRAM1 mRNA expression in parental MCF-7 cells and three ET-resistant counterparts. Data are presented as mean *±* SEM for triplicate samples from one experiment representative of two independent experiments. *P<0.05 by one-way ANOVA. **C.** RT-qPCR of TRAM1 mRNA levels following siRNA-mediated knockdown. Data are presented as mean *±* SEM for triplicates from an experiment representative of three independent experiments. *P<0.05 by One-way ANOVA. **D.** Cell death assessed by Hoechst-PI staining in ET-sensitive and ET-resistant MCF-7 cells after TRAM1 knockdown. Data are presented as mean *±* SEM for triplicate samples from one experiment representative of three independent experiments. *P<0.05 by One-way ANOVA. **E.** RT-qPCR of ISR gene expression following TRAM1 knockdown in ET-sensitive and ET-resistant MCF-7 cells. Data are presented as the mean of triplicate samples from one experiment representative of three independent experiments. Log_2_ fold changes are calculated relative to untransfected control. **F.** Correlation of top ceramide-induced DEGs with the Log_2_ fold changes of the same genes upon TRAM1 knockdown in ET-sensitive and ET-resistant MCF-7 cells. **G.** MCF-7 parental cells stably transfected with a TRAM1-EGFP expression vector or an EGFP-only vector (control). Fluorescent images show GFP localization under each condition. **H.** RT-qPCR of TRAM1 gene expression. Data are presented as mean *±* SEM for triplicate samples from one experiment. **I.** Cell death assessed by Hoechst-PI staining in C8-Ceramide-treated TRAM1-overexpressing and control MCF-7 cells for 48 hours. Values are the mean *±* SEM for triplicate samples from an experiment representative of two independent experiments. *P<0.05 by two-way ANOVA with Geisser-Greenhouse correction. Dose response of MCF-7 is compared to that of each resistant cell line. **J.** Cell death assessed by Hoechst-PI staining in Thapsigargin (Tg)-treated TRAM1-overexpressing MCF-7 cells and control cell lines for 24 hours. Values are mean *±* SEM for triplicate samples from an experiment representative of two independent experiments. *P<0.05 by two-way ANOVA with Geisser-Greenhouse correction. Dose response of MCF-7 is compared to that of each resistant cell line. **K.** Total cell count (Hoechst) and cell death (Hoechst-PI) assessed in parental and TRAM1-overexpressing MCF-7 cells treated with 5 µM of various endocrine therapy drugs (Las: Lasofoxifene, Ela: Elacestrant). Values are mean *±* SEM for triplicate samples from an experiment representative of two independent experiments. *P<0.05 by one-way ANOVA. **L.** Clonogenic growth of parental and TRAM1-overexpressing MCF-7 cells over 14 days in the presence or absence of 1 µM of each endocrine therapy drug. Percent confluence was calculated relative to vehicle-treated controls for each cell line. Values are shown as mean *±* SEM for triplicate samples from an experiment representative of three independent experiments. *P<0.05 by one-way ANOVA.

To further explore the relationship between TRAM1 expression and ceramide sensitivity, we overexpressed TRAM1 in parental MCF-7 cells (**Fig 5G, H**) and observed that TRAM1 overexpression increases sensitivity to both ceramides (**Fig 5I**) and the EnRS-inducer Thapsigargin (**Fig 5J**), indicating that elevated TRAM1 levels may sensitize cells to both EnRS and ceramide-induced cell death. TRAM1 overexpression also improved the survival of parental MCF-7 cells under ET in both short-term (**Fig 5L**) and long-term (**Fig 5M**) conditions.

Together, these findings identify TRAM1 as a potential contributor to the ET-resistant phenotype and suggest that ceramide’s preferential pro-apoptotic effects in ET-resistant cells may be mediated through its interaction with TRAM1.

### Validation of ceramide action in advanced models of ET resistance

Next, we validated our findings using an ET-sensitive patient-derived xenograft organoid (HCI-011) and its fulvestrant-resistant derivative (HCI-011-FR).^64^ We first confirmed the ET-resistant status of HCI-011-FR by observing significant inhibition of organoid growth by fulvestrant and 4-OHT in HCI-011 but not HCI-011-FR PDxOs (**Fig 6A**). Importantly, and consistent with our previous observations, C8-ceramide reduced organoid growth exclusively in the ET-resistant HCI-011-FR, but not in the ET-sensitive HCI-011 (**Fig 6A**). HCI-011-FR also exhibited higher baseline expression of the ISR genes (**Fig S10A**). C8-ceramide treatment induced a stronger activation of ISR genes in HCI-011-FR as compared to HCI-011 (**Fig 6B, C** and **Fig S10B**). Interestingly, the CERK inhibitor NVP-231 also increased ISR gene expression in the HCI-011-FR but not in the parental HCI-011 (**Fig 6D** and **Fig S10C**). Similarly, the PERK activator MK-28 caused a preferential activation of ISR genes in HCI-011-FR (**Fig 6E** and **Fig S10D**) while Thapsigargin activated PERK downstream genes in both the PDxOs (**Fig 6F** and **Fig S10E**). MK-28 and Thapsigargin reduced organoid growth in both HCI-011 and HCI-011-FR, but their effects were more pronounced in HCI-011-FR (**Fig 6G**), thereby validating increased sensitivity of ET resistant models to PERK activation and ceramide-induced preferential activation of PERK in ET-resistant organoids.

**Figure 6:**
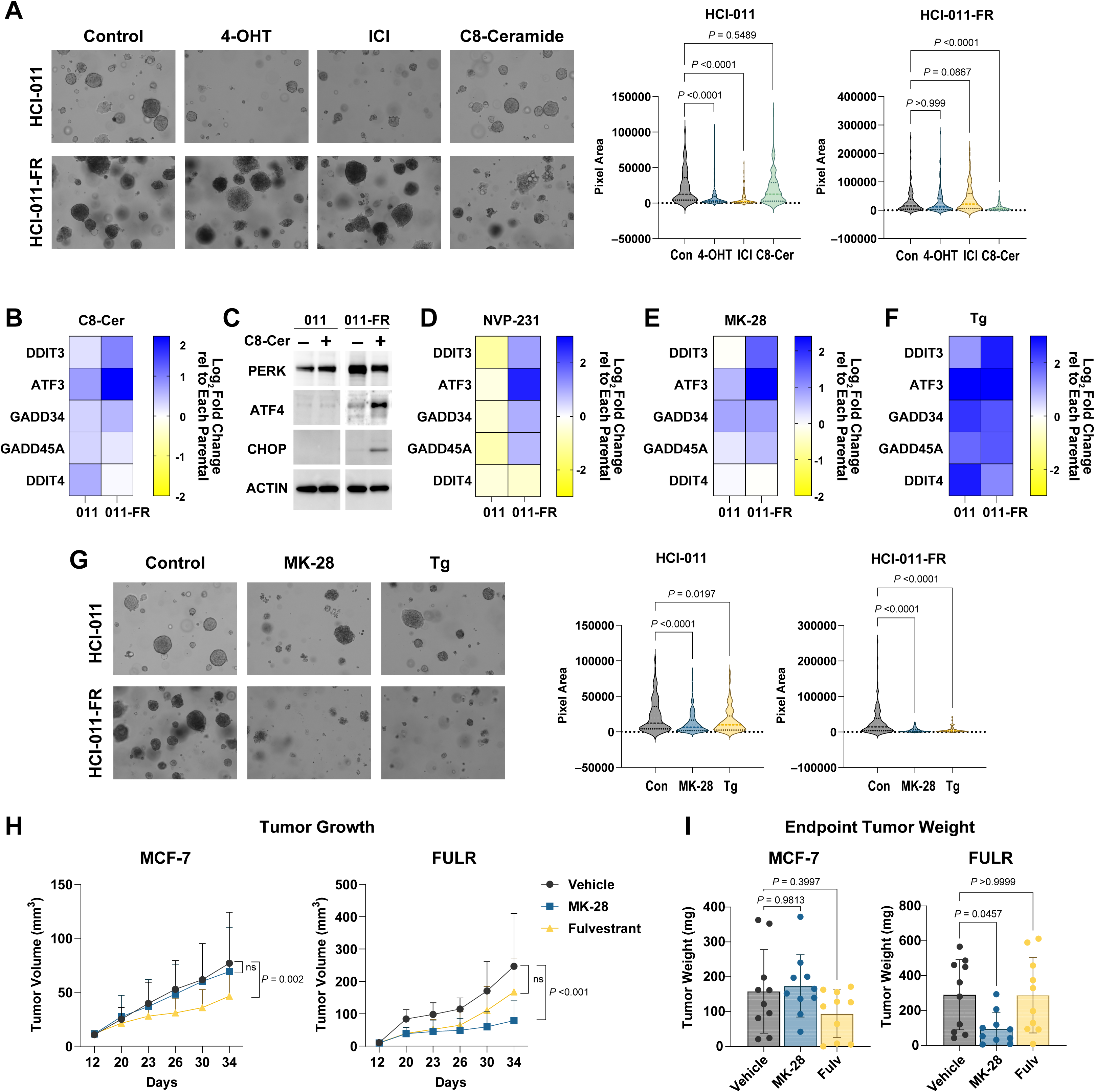
Ceramide action is preserved in advanced preclinical models. A. Growth of ET-sensitive HCI-011 and its fulvestrant-resistant HCI-011-FR patient-derived xenograft organoids (PDxO) treated with vehicle, 1 µM 4-OHT, 1 µM ICI, or 10 µM C8-Ceramide for 7 days. Data are presented as violin plots representing the distribution of organoid sizes under each condition. *P<0.05 by one-way ANOVA. B. RT-qPCR of ISR genes from HCI-011 and HCI-011-FR treated with or without 10 µM C8-Ceramide for 24 hours. Data represent the mean of triplicate samples from one experiment, shown as Log_2_ fold change relative to vehicle-treated controls. C. Western Blots on HCI-011 and HCI-011-FR PDxO treated with or without 10 µM C8-Ceramide for 24 hours. Representative blots from two independent experiments are shown. D. RT-qPCR from HCI-011 and HCI-011-FR PDxO treated with 10 µM NVP-231 for 24 hours. Means of triplicate samples are shown from one experiment as Log_2_ fold change relative to vehicle-treated controls in each model. E. RT-qPCR from HCI-011 and HCI-011-FR treated with or without 10 µM MK-28. Means of triplicate samples are shown from one experiment as Log_2_ fold change relative to vehicle-treated controls in each model. F. RT-qPCR from HCI-011 and HCI-011-FR treated with or without 1 µM Thapsigargin. Means of triplicate samples are shown from one experiment as Log_2_ fold change relative to vehicle-treated controls in each model. G. Growth of HCI-011 and HCI-011-FR PDxO treated with vehicle, 10 µM MK-28, or 1 µM Thapsigargin for 7 days. Data are presented as violin plots representing the distribution of PDxO sizes under each condition. *P<0.05 by one-way ANOVA. H. Growth of xenograft tumors derived from ET-sensitive MCF-7 and fulvestrant-resistant MCF-7-FULR cells in nude mice. Mice were treated with vehicle, fulvestrant, or MK-28 (5 animals per group, 2 tumors per animal) for 3 weeks. Data represent mean *±* SD for 10 tumors per group. *P<0.05 by two-way ANOVA with Geisser-Greenhouse correction. I. Endpoint tumor weights from MCF-7 and MCF-7-FULR xenografts in nude mice treated with vehicle, fulvestrant, or MK-28. Data represent mean *±* SD for 10 tumors per group. *P<0.05 by one-way ANOVA.

While delivery of ceramides as a potential cancer treatment has been explored,^20^ in general this approach has been challenging. As such, we sought to determine whether ET-resistant cells retained their sensitivity to the PERK activator MK-28 when grown as xenograft tumors *in vivo*. We grew MCF-7 parental and MCF-7-FULR xenograft tumors in the mammary fat pad of female nude mice, where we observed moderate sensitivity to Fulvestrant in MCF-7 parental tumors (**Fig 6H, I**). Importantly, we found that treatment with MK-28 significantly reduces MCF-7-FULR tumor growth while having no effect on the growth of MCF-7 tumors (**Fig 6H, I**). These results suggest that ET-resistant cells retain their sensitivity to PERK activation *in vivo* and suggest that this approach may be useful to treat ET-resistant tumors.

## DISCUSSION

Our previous studies have established that ET-resistant breast cancer models exhibit a characteristic reduction in ceramide levels, that may represent an adaptation essential for the survival of ET-resistant cells. Here, we sought to delineate the molecular basis of this dependency on diminished ceramide abundance. Our findings identify the EnR membrane protein TRAM1 as a potential mediator of ceramide action in ET resistance. We demonstrate that TRAM1, known to attenuate EnRS, is enriched in ET-resistant cells, where it plays a critical role in maintaining their viability. Our data suggest that increased TRAM1 sensitizes cells to ceramides, which may function to disrupt TRAM1 action and induce unmitigated EnRS and preferential cell death in ET-resistant models.

EnRS is increasingly recognized as a key adaptive mechanism during tumor onset, progression, and therapy resistance.^41, 65–67^ In luminal breast cancer, ET has been shown to induce EnRS, promoting autophagy *via* AMPK activation and supporting cellular survival.^42, 43^ Long-term ET has been associated with sustained EnRS,^44, 45^ and previous studies demonstrated that this long-term adaptation to EnRS can be exploited to cause further EnRS through knockdown of the EnR chaperone GRP78 to restore tamoxifen-sensitivity in ET-refractory breast cancer cells.^46^ Consistent with this, we show that pharmacologic activation of PERK selectively induces cytotoxicity in multiple ET-resistant models, likely due to elevated baseline EnRS levels. Provision of exogenous ceramides exploits this vulnerability by activating PERK signaling, thereby driving preferential cytotoxicity in ET-resistant cells. Although increased sensitivity to EnRS and PERK activation was consistently observed across multiple ET-resistant models, there were also differences observed in terms of response to EnRS and PERK activation in different models of ET resistance, suggesting EnRS can engage distinct downstream pathways depending on the resistance context. The model-specific responses to EnRS and the differential effects of ceramides across ET-resistant models were not addressed in the present study and represent avenues for future investigation.

Ceramides, as bioactive lipids, are known to interact with diverse proteins, thereby influencing key cellular processes. Our proteomic analysis identified several ceramide-interacting proteins (CIPs) with established roles in ET resistance, including *PRKCA*, *ERBB2*, and *IKBKB*, as well as several additional CIPs whose connection to ET resistance remains unknown. Although various ceramide-binding motifs have been proposed the functional consequences of these interactions remain incompletely understood. Given the heterogeneity of ceramide responses across ET-resistant models, it is plausible that ceramide-CIP interactions vary between different resistance subtypes. An in-depth understanding of the biological roles of ceramide-binding motifs is therefore needed. Notably, TRAM1 has previously been identified as a ceramide-interacting protein;^54^ however, our study is the first to establish its functional relevance in the context of ET resistance. The mechanism underlying TRAM1 upregulation in ET resistance remains to be fully elucidated. *TRAM1* is located at the cytogenetic location 8q13.3. While ceramides have shown promise *in vitro*, their therapeutic application presents considerable challenges.

Since their discovery as apoptosis inducers, attempts to deliver ceramides clinically have been hindered by significant formulation and delivery obstacles (Clinical Trials: NCT02834611, NCT00008320).^20^ While direct ceramide administration is a challenging clinical strategy, understanding the molecular basis of increased ceramide sensitivity in ET-resistant breast cancer cells reveals new therapeutic opportunities. Increasing endogenous ceramide levels through CERK inhibition may offer one such avenue. Additionally, emerging targets such as TRAM1 present potential pharmacological targets for therapeutic developments. Finally, the selective sensitivity of ET-resistant cells to PERK activation could be exploited using pharmacologic PERK activators, offering a novel strategy for targeting ET-resistant luminal breast cancer in the clinic.

## METHODS

### Materials

Lipid standards used in LC-MS analysis were purchased from Avantipolar Lipids (AL, USA): C17 ceramide (#860517), C17 sphingomyelin (#860585), and C17 glucosylceramide (#860569). LC-MS columns and guard cartridges were purchased from Phenomenex (Torrance, CA, USA). NVP-231 (#13858), C8 ceramide (d18:1/8:0) (#62540), Lasofoxifen (#21754), Elacestrant (#37298), Thapsigargin (#10522),

Tauroursodeoxycholic acid (20277), 4-phenylbutyric acid (#11323), Salubrinal (#14735), and ISRIB (#41879) were purchased from Cayman Chemical Company (MI, USA). GSK2616157 (HY-13820), MK-28 (HY-137207) and Fulvestrant (HY-13636) were purchased from MedChemExpress (NJ, USA). siTRAM1 (AM16708) and Silencer^TM^ negative control (siNEG) (Thermo Fisher Scientific, Waltham, MA, USA) were purchased from Thermo Fisher Scientific. 4-hydroxytamoxifen (4-OHT) was purchased from Sigma (#H7904). Primary antibodies to PERK (#3192), ATF4 (#11815) and CHOP (#2895) were purchased from Cell Signaling Technology and β-actin (mouse) primary antibody was purchased from Sigma (#A5441). Goat anti-rabbit (#31460) and anti-mouse (#31430) HRP-conjugated secondary antibodies were purchased from Invitrogen (Waltham, MA, USA). pacFA Ceramide (#90040P) and C16 ceramide (d18:1/16:0) (#860516P) were purchased from Avantipolar Lipids (AL, USA).

### In Vitro Culture

ET-sensitive MCF-7 and its ET-resistant derivative MCF-7-HER2, MCF-7-TAMR and MCF-7-FULR were generously provided by Dr. Rachel Schiff (Baylor). ET-sensitive T47D cell and its ET-resistant derivatives T47D/PKCα and T47D:C42-ED were generously provided by Dr. Debra Tonetti (UIC). All cell lines were maintained in their standard growth conditions as previously described.^35, 36, 41, 68–70^ Cell line authentication using short tandem repeat (STR) and mycoplasma contamination testing were performed for all cell lines. All drug treatments were performed in respective growth media of each cell line unless otherwise specified.

Patient-derived xenograft organoids (PDxO) were kindly provided by Dr. Alana Welm (University of Utah, Huntsman Cancer Institute) and were cultured and maintained as previously described.^41^ Briefly, PDxO embedded in 100% matrigel (Corning) were cultured in Advanced DMEM/F12 supplemented with 5% FBS, 1% Glutamax (Gibco, #35050-061), 0.1% hydrocortisone (Sigma, #H0135), 0.1% Gentamycin (Sigma, #345815), and 0.01% hEGF (Gibco, #PHG0311) with the following additives: 10 µM Y-27632 (SelleckChem #S1049), 1 mM N-acetyl cysteine (Sigma, #A7250), 100 ng/mL basic FGF2 (Thermo #68-8785-82), and 10 nM estradiol (E2) (Sigma #E2758).

### Total RNA-seq

ET-sensitive MCF-7 cells and three of its ET-resistant derivatives (MCF-7-HER2, MCF-7-TAMR, and MCF-7-FULR) were treated with 10 µM C8-ceramide or ethanol vehicle control in respective growth media (see above). After 24h of treatment, RNA was isolated from the cells using Trizol reagent (Thermo Fisher). RNA samples were sent to Genomics Research Core at University of Illinois Chicago for human mRNA sequencing. Sequencing was carried out on NovaSeq 6000 (Illumina), S4 flowcell, 2/150 bp, and 60 paired-end reads per sample. Sequencing libraries for Illumina sequencing were prepared in one batch in 96-well plate using CORALL Total RNA-seq Library Prep Kit with Unique Dual Indices with RiboCop HMRv2 rRNA Depletion Kit (Lexogen).

### Functional Characterization of Differentially Expressed Genes

The Database for Annotation, Visualization and Integrated Discovery (DAVID) (https://davidbioinformatics.nih.gov/) was used for functional annotation of the DEGs. Functional annotation was performed using Gene Ontology (GO) terms (biological process, molecular function, and cellular component), KEGG pathways, and Reactome pathways. A modified Fisher exact test was used to calculate the enrichment P-values. Benjamini-Hochberg correction was applied to adjust for multiple testing, and a P-value threshold of < 0.05 was used to identify significantly enriched terms and pathways.

Gene Set Enrichment Analysis (GSEA) (version 4.3.2)^71^ was performed using the GSEA software from the Broad Institute (UCSD). List of DEGs, based on log2 fold change was used as input for GSEA. The gene sets used for enrichment analysis were Molecular Signatures Database (MSigDB) h.all.v2024.1. The analysis was run with 1000 permutations, phenotype taken as permutation type, enrichment statistic was weighted and genes were ranked according to Signal2Noise ratio. A false discovery rate (FDR) corrected P-value (q-value) of < 0.1 and a normalized enrichment score (NES) of *|*NES*|* > 2 were used as significance thresholds to identify significantly enriched gene sets. All Venn diagrams were prepared using Venny 2.1 (https://bioinfogp.cnb.csic.es/tools/venny/).

### Proteomic Analyses

Cell pellets were collected from MCF-7 and HER2 cell lines. The pellets were resuspended in 400 µL of cold PBS containing protease inhibitors, along with 100 µL of a 20% SDS solution. Sonication was performed on ice (40% amplitude, 10s on, 5s off x3). Protein concentrations were measured, and samples were normalized to ensure 1 mg of protein per sample. 5 µM C16-PacCeramide was added to each lysate. The samples were incubated at 37°C for 15 minutes in the dark. After the incubation, samples were irradiated at 365 nm for 30 minutes. The samples were then placed on ice and subjected to protein precipitation using 500 µL of ice-cold methanol and 200 µL of ice-cold chloroform. The samples were vortexed and centrifuged at maximum speed (16,900 rcf) for 10 minutes at 4°C. The liquid layers were carefully removed, and 1x volume of methanol was added to the protein pellet, which was then sonicated and centrifuged for 5 minutes at maximum speed at 4°C. The supernatant was discarded, and the pellets were resuspended in 4% SDS in 1xPBS with sonication.

For the click reaction, a master mix was prepared containing CuSO4 (1 mM), TBTA (250 µM), and biotin-PEG3-azide (250 µM). Freshly prepared TCEP (1 mM, pH 7) was added to the samples, followed by the addition of the master mix. The samples were vortexed and incubated at 37°C for 1 hour. After the reaction, protein precipitation was repeated as described above to remove excess reagents. After precipitation, the pellets were dissolved in 2% SDS in PBS using sonication. Aliquots were taken for protein measurement via the BCA assay. Protein concentrations were normalized across samples. For proteomics, approximately 900 µg of protein from each sample was transferred to new microcentrifuge tubes for enrichment. High-capacity neutravidin-agarose resin was resuspended, and 60 µL bead slurry was added to each sample. The beads were washed three times with 0.2% SDS in PBS, followed by centrifugation at 2000 rcf for 2 minutes at room temperature. The beads were resuspended in 400 µL 1% Brij solution and incubated with the samples for 1.5 hours at room temperature on a rotator. After incubation, the beads were centrifuged (2 minutes, 2000 rcf, room temperature), and the supernatant was carefully removed. The beads were washed three times with 0.2% SDS in 1xPBS and three more times with PBS. After final centrifugation and removal of the supernatant, the enriched beads were stored at −80°C for subsequent proteomics analysis. Protein digestion on the beads, sample prep for proteomics and analysis were performed as previously described.^72^

### STRING-based functional analysis of CIPs

Functional enrichment analysis was performed using the STRING database’s (https://string-db.org/)^73^ built-in analysis tools. Gene Ontology (GO) terms (biological process, molecular function, and cellular component), KEGG pathways, and Reactome pathways were analyzed. A false discovery rate (FDR) correction was applied using the Benjamini-Hochberg method to adjust for multiple testing. A significance threshold of P < 0.05 (FDR-corrected) was used to identify enriched terms and pathways.

### Generation of TRAM1-overexpressing cell lines

pTRAM1-EGFP was a gift from Michele Pagano (Addgene plasmid #210365: http://n2t.net/addgene:210365; RRID: Addgene 210365).^74^ pEGFP-N1-1x was a gift from Antony K. Chen (Addgene plasmid #172281: http://n2t.net/addgene:172281; RRID: Addgene 172281).^75^ Lentiviruses were generated with transfection of plasmids (10 µg of pLenti-TRAM1, 5 µg of pDNRF, and 3 µg of pVSVg) in HEK293T cells with polyethylenimine (PEI). After overnight, the media changed to DMEM with non-essential amino acids. Lentiviruses were collected and sterilized with a 0.45um syringe filter after 2 days. Parental ET-sensitive cells were infected with the lentiviral particles with regular growth media at 50:50 v/v for 24 hours and selected with puromycin for 72 hours.

### Gene Knockdown Studies

Cells were transfected with siRNA targeting TRAM1 (s23890) or a non-targeting control (siNeg) (Thermo Fisher Scientific, Waltham, MA, USA) using DharmaFECT 1 (Dharmacon, Lafayette, CO, USA). Transient transfection was performed with a final concentration of 20 nM of siRNA in 4% DharmaFECT in OptiMEM (Gibco, Waltham, MA, USA). Media was changed to regular media at 24-h post-transfection, and endpoint assays were performed at 72-h post-transfection.

### Cell Viability Assay

Cell confluency and cell death were analyzed with a Celigo Imaging Cytometer (Nexcelom Bioscience, Lawrence, MA, USA), as previously described.^19^ Briefly, cells were treated with Hoechst 33,432 (Life Technologies, Carlsbad, CA, USA) and propidium iodide (PI) (final concentration 1 µg/mL) for 30 min and incubated for 30 min at 37°C. PI-positive cells and total cells were then detected in the red and blue fluorescence channels, respectively. The percentage of dead cells was determined as the number of dead cells (positive red fluorescence signal) over the total number of cells (positive blue fluorescence signal) per well.

### Clonogenic Assay

Cells were seeded in 6-well plate in 1,000 cells per well density in growth media (see above) and treated with ET at final concentrations of 1 µM. Media with treatment was refreshed every 3–4 days for 2 weeks. After 2 weeks, plates were scanned with a Celigo Imaging Cytometer (Nexcelom Bioscience). Confluence was calculated from endpoint Hoechst staining using the confluence application.

### RT-qPCR

Total RNA isolation with Trizol (Ambion, Austin, TX, USA), cDNA synthesis, and RT-qPCR were performed as previously described with 36B4 and GAPDH as housekeeping controls.^19^ Primer sequences are listed in **Table S8**.

### Western Blot

Whole-cell extracts were prepared using M-PER reagent (Thermo Scientific, 78503 Waltham, MA, USA). Proteins were denatured and then separated by NovexTM WedgeWellTM 4–12% Tris-Glycine gel (Invitrogen, XP04120) and transferred to nitrocellulose membrane (Thermo Scientific, 88018). The membrane was blocked with 5% non-fat dry milk in 1% TBST and incubated overnight with primary antibodies (diluted in 5% BSA in 1% TBST) at 4 °C. Membranes were then washed and incubated with HRP-conjugated-secondary antibodies. The signal was visualized using Chemidoc MP (Bio-Rad laboratories, Hercules, CA, USA) with the Pierce Supersignal West Pico Chemiluminescent substrate (Thermo Scientific).

### PDxO Growth Assessment

Patient-derived xenograft Organoids (PDxO) were grown as previously described.^41^ For growth assessment, 500 organoids were seeded per well in a 24-well plate (Corning) in a 20 µl matrigel (Corning) dome on a 5 µl base layer of matrigel (n=3 for each treatment group). Organoids were grown for 5 days in their growth media and treatment was initiated on day 6 which continued for another 7 days. Media was changed every 3 days along the entire length of the experiment. All drugs were prepared in the growth media of the organoids. Images were acquired on 3 random fields of view per well on days 0, 3, and 7 of the treatment at 20X magnification of EVOS M5000 (Invitrogen) microscope. All images were blinded and organoid area was calculated in ImageJ. Organoid growth was estimated by the distribution of organoid area under each treatment condition.

### Animal Studies

All mouse procedures were approved by the University of Illinois Chicago Animal Care Committee (ACC). 5 *×* 10^6^ MCF-7 parental and MCF-7-FULR cells were injected into the fat pad of the #4 mammary glands of 6 to 8-week old female athymic nude-foxn1*^null^* mice. Cell suspensions were injected in a 0.5 cc growth factor reduced matrigel (Corning). Estrogen (E2) was administered *via* oral route through drinking water at a final concentration of 0.0008%. Once tumors became palpable, drugs were administered in the following way: (a) Fulvestrant was administered once a week at the dose 5 mg/kg body weight of mice in a 1:9 ethanol:peanut oil vehicle (final volume of injection: 0.1 cc) with a 27G needle as a subcutaneous injection, (b) MK-28 was dissolved in 50% glycerol and 15% DMSO in 1x PBS and administered three times per week as intraperitoneal injection at 10 mg/kg body weight of mice. Tumor size was measured by digital caliper over time and tumor volume was calculated using the formula: 0.5 *×* (width)^2^ *×* (length). Mice were euthanized according to institutional guidelines and endpoint tumor weight was recorded.

### Mining and Analysis of Patient Tumor Data

Primary human breast tumor (METABRIC) dataset^76^ was accessed through cBioPortal. Gene expression data and corresponding clinical data, including relapse-free survival and PAM50 subtype classifications were analyzed for luminal breast cancer patients on ET using the in-built online analyses tools on cBioportal. Kaplan-Meier Plotter (https://kmplot.com/) was utilized to investigate prognostic significance of specific gene(s) in luminal breast cancer patients. Median expression level was used to sort patients into high and low expression groups. Hazard ratio (HR), 95% confidence intervals (CI), and P-values were extracted from the generated plots. Log-rank test was used to assess the statistical significance of differences in survival between the stratified groups.

### Statistical Analyses

All data are represented as mean +/- SEM from at least 3 independent samples unless otherwise specified. Statistical analyses were performed using GraphPad Prism v10.0. Student’s t-test and one-way or two-way ANOVA were used as appropriate (as mentioned in individual figure legends). All P-values were two-sided and statistically significant change was considered for P<0.05.

## DATA AVAILABILITY

All data generated or analysed during this study are included in this article (and its supplementary information files). The datasets generated during and/or analysed during the current study are available from the corresponding author(s) on reasonable request.

## SUPPLEMENTARY DATA LEGENDS

### FIGURES

**Figure S1.**
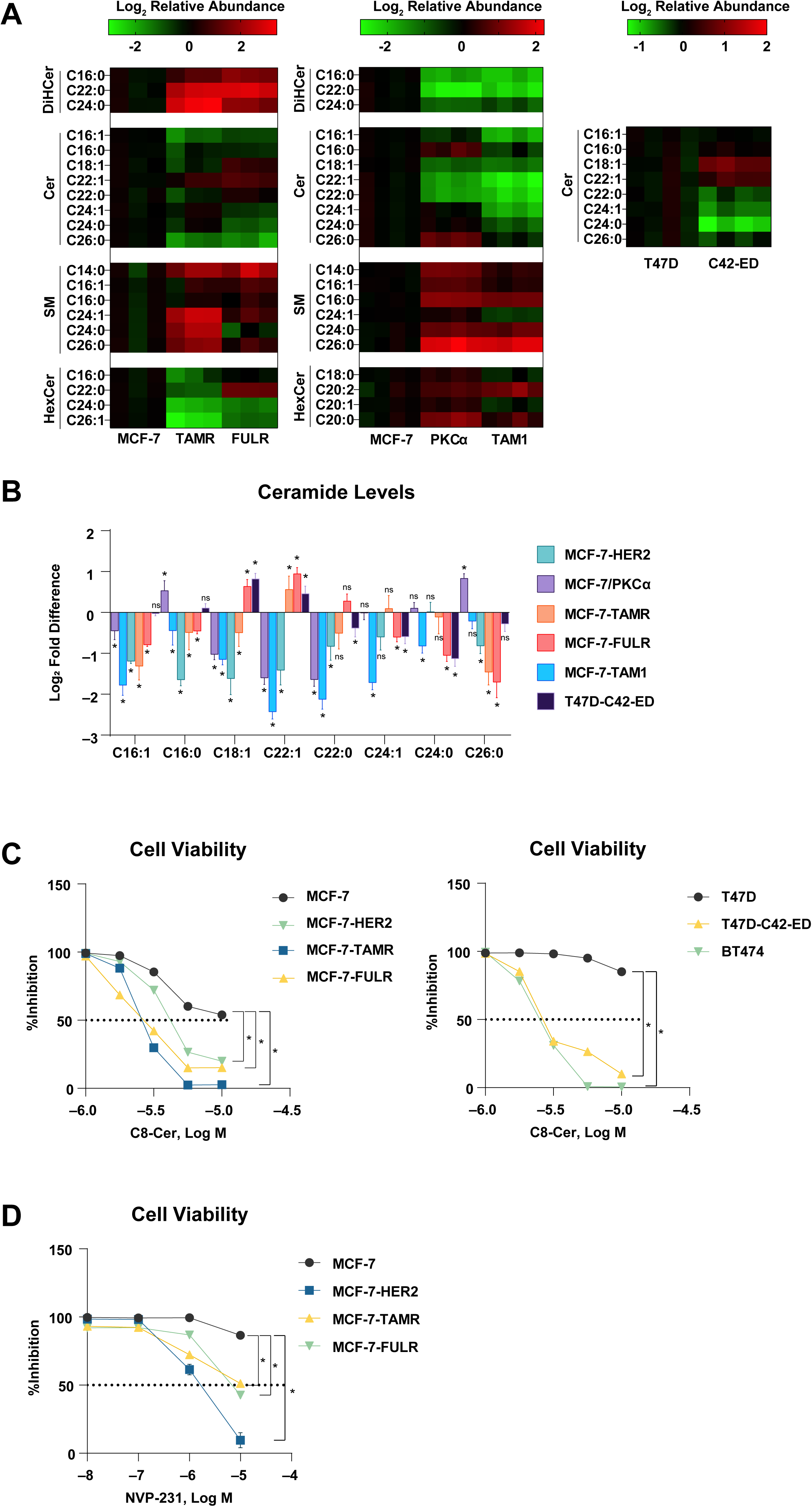
Sphingolipids in multiple ET-resistant cell lines. **A.** Sphingolipid abundance in multiple ET-resistant cells. Relative abundance of total ceramides (Cer), dihydroceramides (DiHCer), hexosylceramides (HexCer), and sphingomyelins (SM) in two different sets of parental versus ET-resistant MCF-7 cell lines, along with ceramide levels in parental T47D versus long-term estrogen depleted derivative (T47D-C42-ED). **B.** Summary of ceramide levels in multiple ET-resistant models. The level of each ceramide in ET-resistant models is reported as fold difference relative to their respective parental ET-sensitive cell line. Data are presented as mean *±* SEM for triplicate samples from one experiment. *P<0.05 by unpaired T-test with Welch’s correction compared to respective parental cells. **C.** Cell death assessed from Hoechst-PI endpoint staining in a panel of ET-sensitive and ET-resistant breast cancer cells treated with increasing concentrations of C8-ceramide for 48 hours. Percent reduction in the number of live cells is shown. Data are presented as mean *±* SEM for triplicate samples from one experiment representative of more than three independent experiments. *P<0.05 by two-way ANOVA with Geisser-Greenhouse correction. **D.** Cell death assessed from Hoechst-PI staining in breast cancer cells treated with increasing concentrations of NVP-231 for 48 hours. Data are presented as mean *±* SEM for triplicate samples from one experiment representative of more than three independent experiments. *P<0.05 by two-way ANOVA with Geisser-Greenhouse correction.

**Figure S2.**
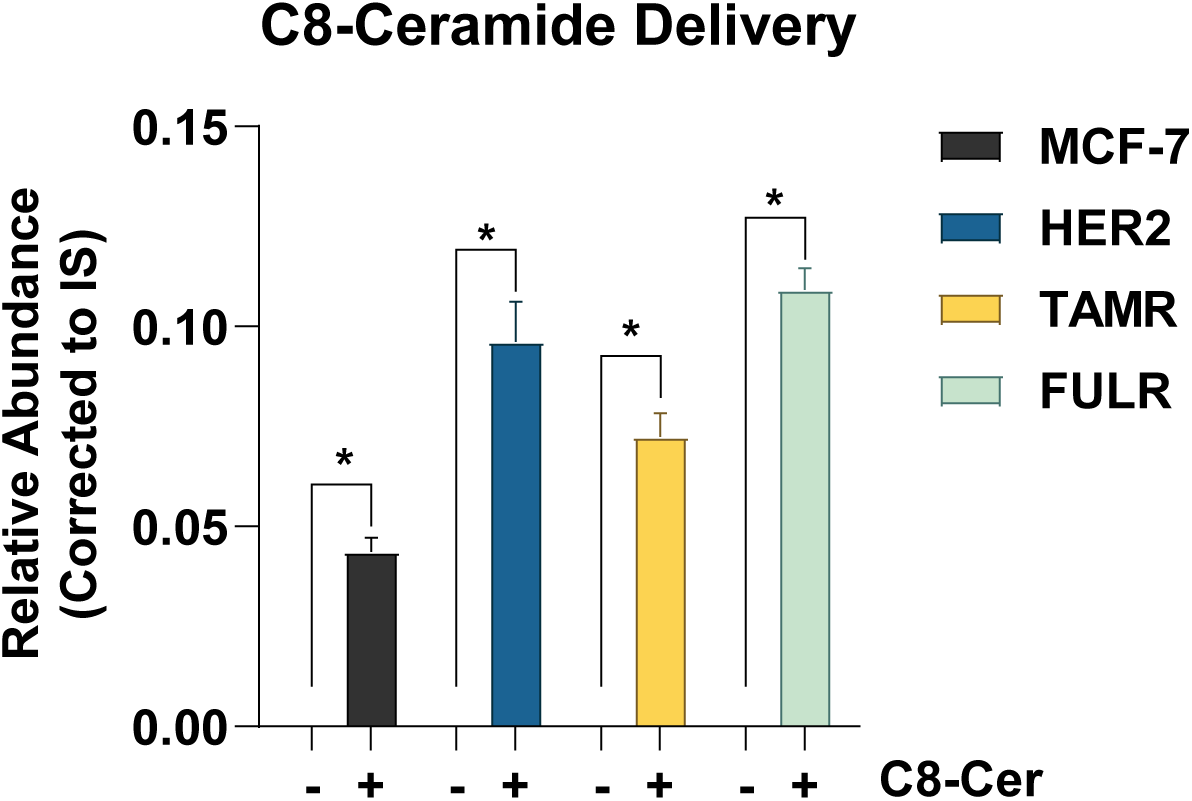
Intracellular delivery of C8-ceramide. Endogenous levels of C8-Ceramide measured in ET-sensitive and ET-resistant MCF-7 cells treated with 10 µM C8-Ceramide for 24 hours. Data are presented as mean *±* SEM for triplicate samples from one experiment. *P<0.05 by unpaired Student’s t-test with Welch’s correction comparing means of C8-ceramide treated groups to their respective vehicle-treated controls.

**Figure S3.**
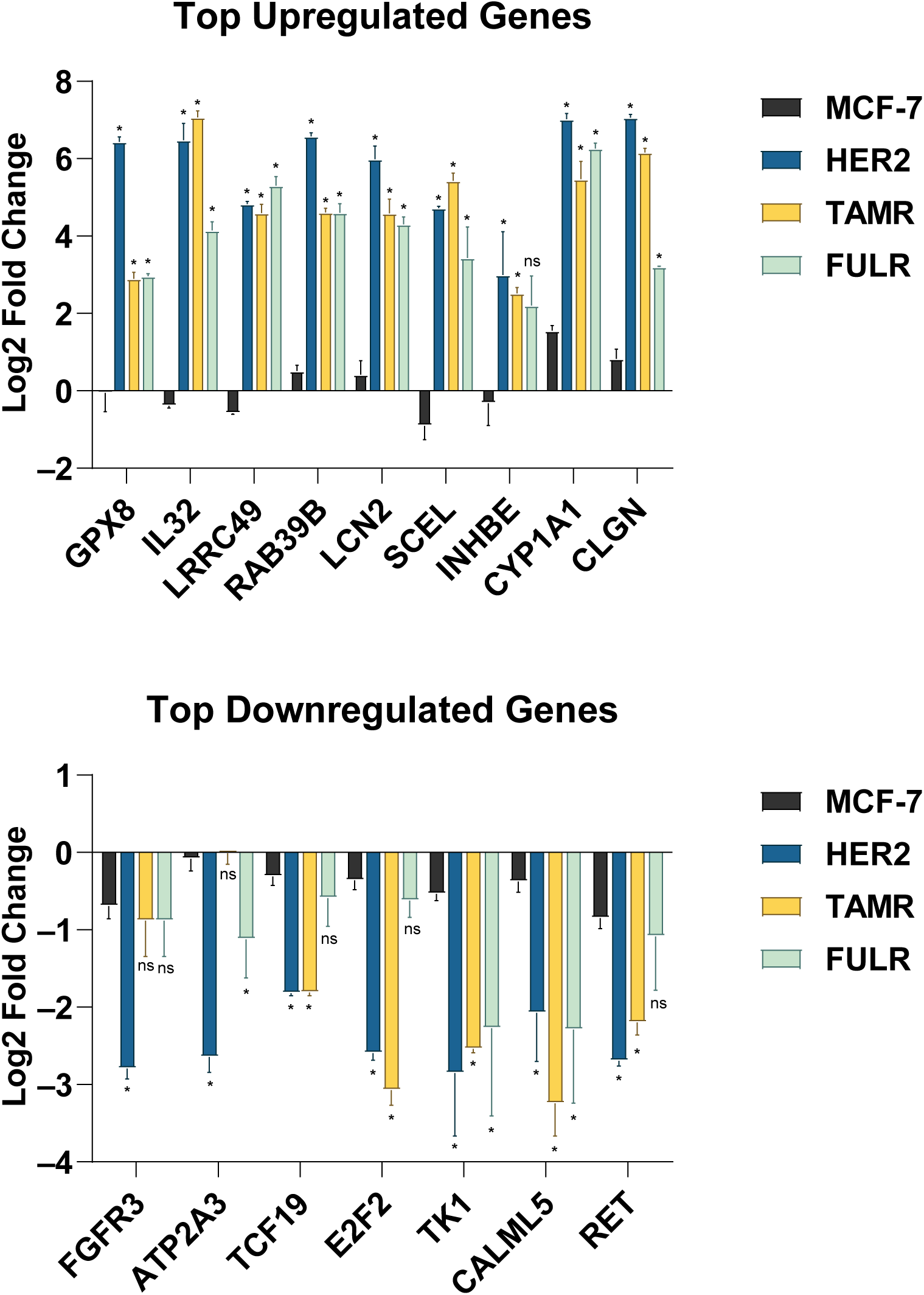
Validation of RNA-seq by RT-qPCR. RT-qPCR of top ceramide-induced genes that are differentially expressed in all ET-resistant cells but not in the ET-sensitive parental cells. Data are presented as mean *±* SEM for triplicate samples from one experiment. *P<0.05 by one-way ANOVA for each gene.

**Figure S4.**
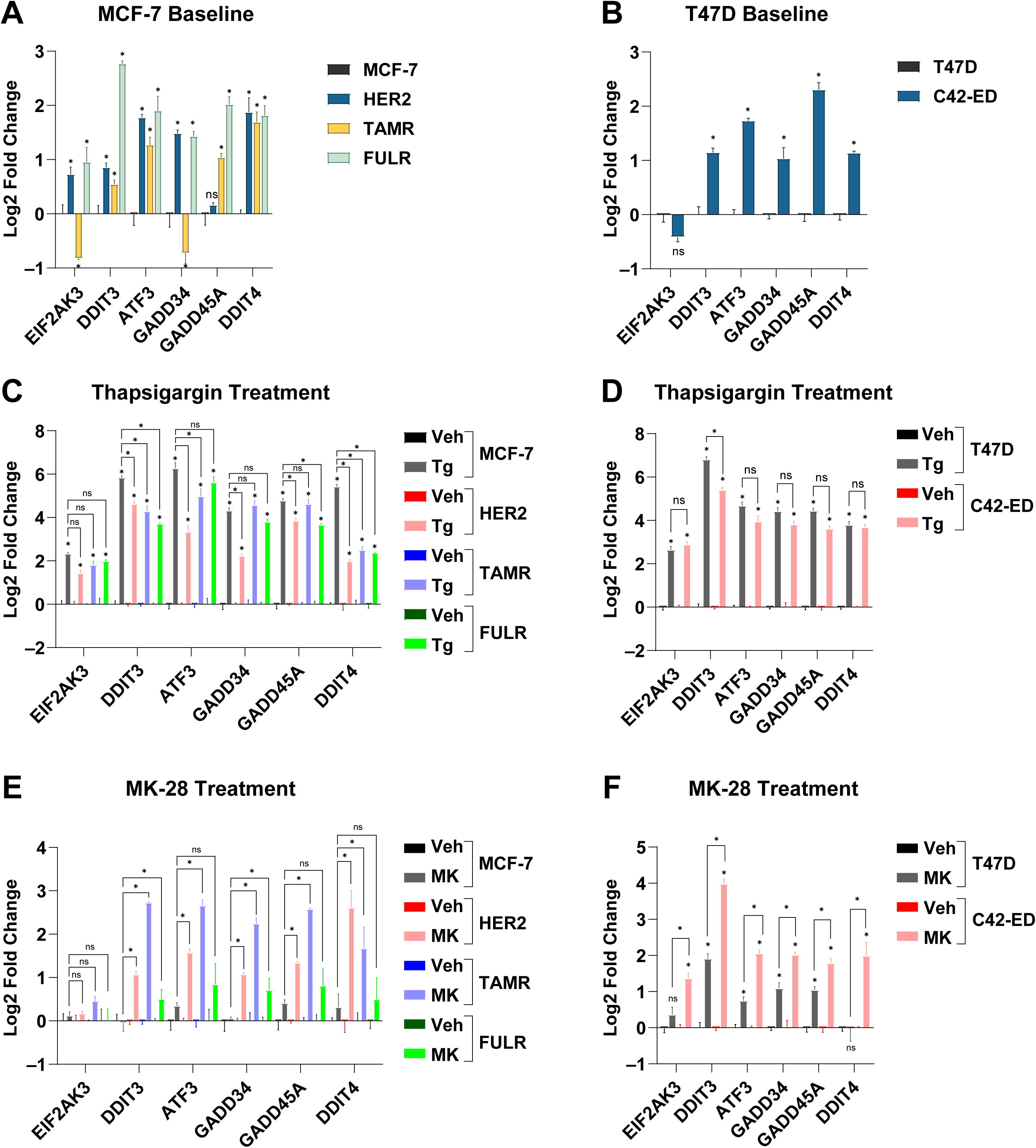
ISR gene expression in ET-sensitive and ET-resistant cells. **A.** RT-qPCR of ISR genes in ET-resistant MCF-7 cells relative to their parental ET-sensitive counterpart. Data are presented as mean *±* SEM for triplicate samples from an experiment representative of three independent experiments. *P<0.05 by one-way ANOVA. **B.** RT-qPCR of ISR genes in ET-resistant T47D cells relative to their parental ET-sensitive counterpart. Data are presented as mean *±* SEM for triplicate samples from one experiment. *P<0.05 by unpaired Student’s t-test with Welch’s correction. **C.** RT-qPCR following 24 hours of 1 µM Thapsigargin treatment in ET-resistant MCF-7 cells. Data are presented as mean *±* SEM for triplicate samples from an experiment representative of two independent experiments. *P<0.05 by one-way ANOVA. **D.** RT-qPCR following 24 hours of 1 µM Thapsigargin treatment in ET-resistant T47D cells. Data are presented as mean *±* SEM for triplicate samples from one experiment. *P<0.05 by unpaired T-test with Welch’s correction. **E.** RT-qPCR following 24 hours of 10 µM MK-28 treatment in ET-resistant MCF-7 cells. Data are presented as mean *±* SEM for triplicate samples from an experiment representative of two independent experiments. *P<0.05 by one-way ANOVA. **F.** RT-qPCR following 24 hours of 10 µM MK-28 treatment in ET-resistant T47D cells. Data are presented as mean *±* SEM for triplicate samples from one experiment. *P<0.05 by unpaired T-test with Welch’s correction.

**Figure S5.**
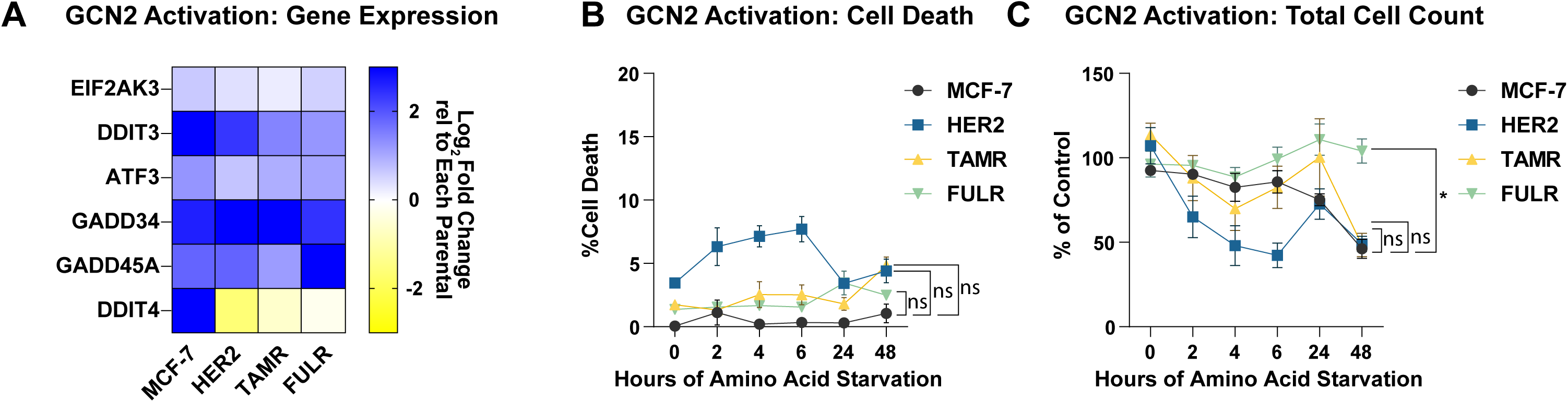
Amino acid starvation of ET-sensitive and -resistant MCF-7 cells. **A.** RT-qPCR from ET-sensitive and ET-resistant MCF-7 cells cultured in media with or without amino acids for 4 hours to activate GCN2, another upstream activator of ISR. Heatmap shows means of triplicates from one experiment. **B-C.** Total cell number assessed by Hoechst (**B**), and cell death assessed by Hoechst-PI staining (**C**) in ET-sensitive and ET-resistant MCF-7 cells cultured in media with or without amino acids for up to 48 hours. Data are presented as mean *±* SEM for triplicates from one experiment representative of two independent experiments. P>0.05 by two-way ANOVA with Geisser-Greenhouse correction.

**Figure S6.**
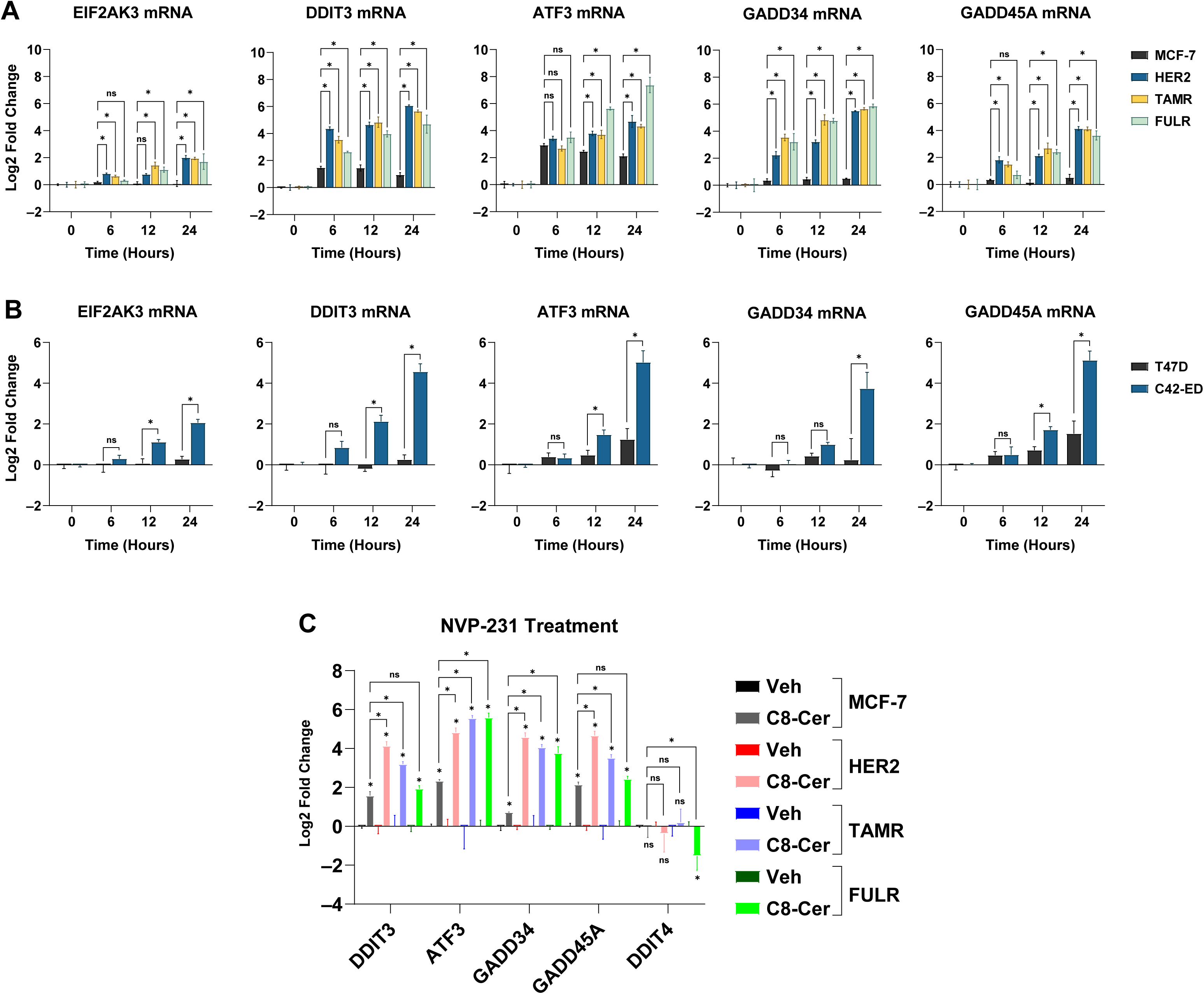
Expression of ISR genes in ET-sensitive and ET-resistant cells from elevated ceramide levels. **A.** RT-qPCR showing gene expression over 24 hours of 10 µM C8-ceramide treatment in MCF-7 cells. Data are presented as mean *±* SEM, with log_2_ fold changes calculated relative to 0-hour controls. *P<0.05 by one-way ANOVA. **B.** Same as **A**, in T47D cells. *P<0.05 by unpaired T-test with Welch’s correction. **C.** RT-qPCR showing gene expression following 10 µM NVP-231 treatment for 24 hours in MCF-7 cells. Data are presented as mean *±* SEM of triplicate samples. *P<0.05 by one-way ANOVA.

**Figure S7.**
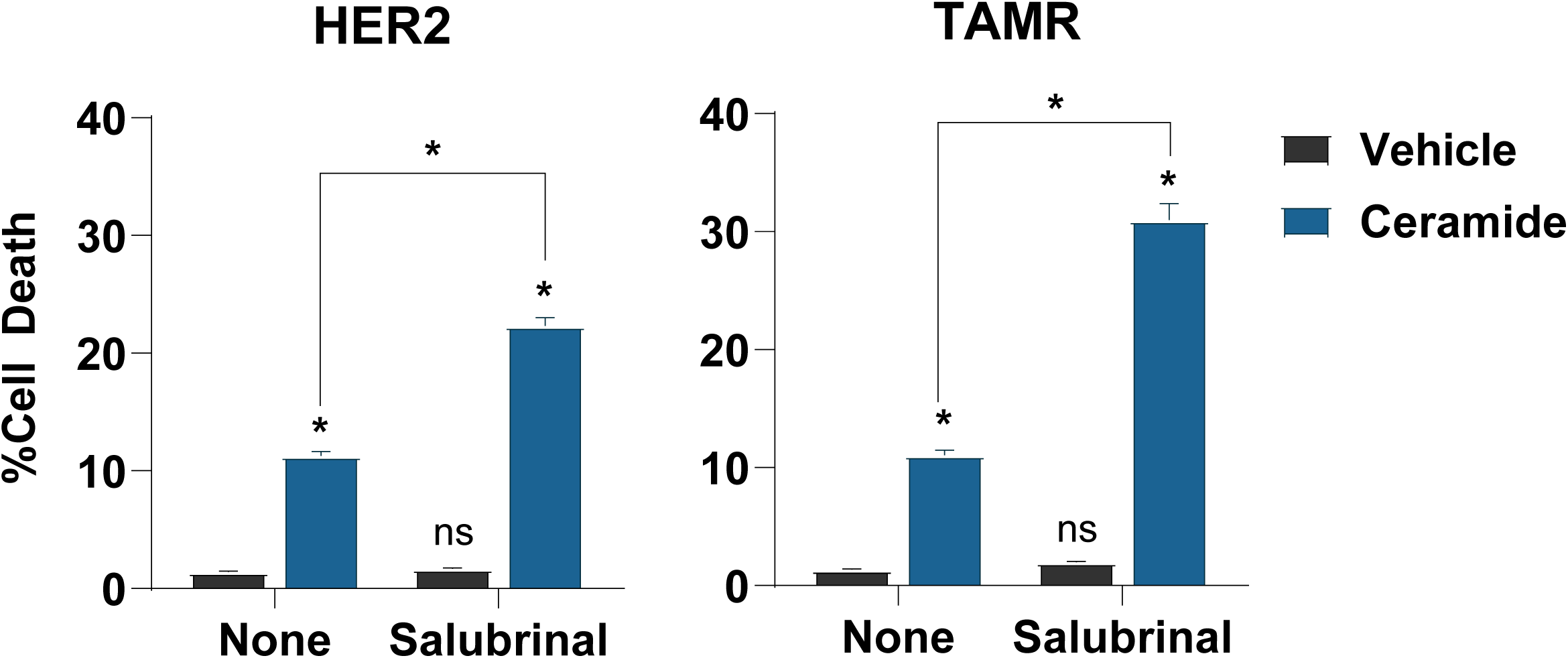
Salubrinal co-treatment with C8-ceramide to ET-resistant MCF-7 cells. Cell death assessed by Hoechst-PI staining in MCF-7-HER2 and MCF-7-TAMR cells following the treatment of 10 µM Salubrinal along with 5 µM C8-Ceramide for 24 hours. Data are presented as mean *±* SEM for triplicate samples from one experiment. *P<0.05 by two-way ANOVA with Geisser-Greenhouse correction.

**Figure S8.**
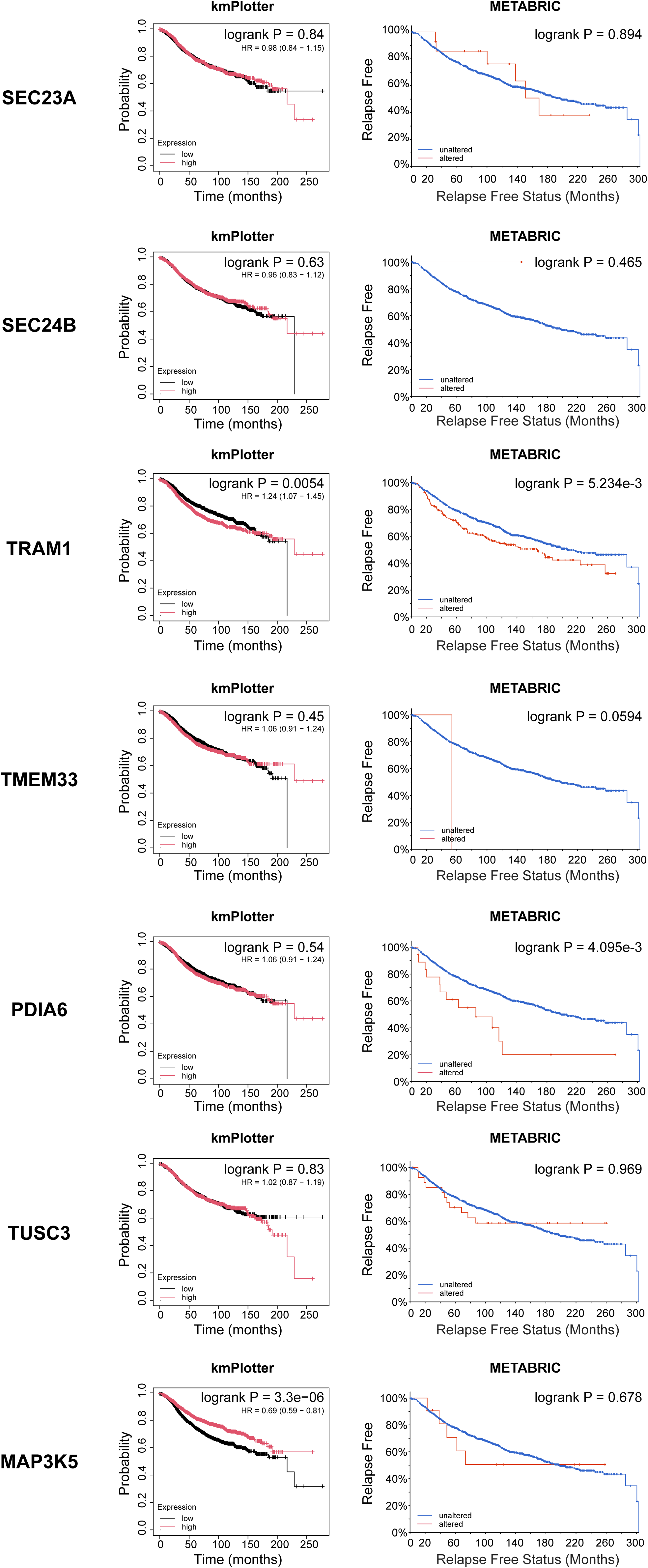
Kaplan-Meier plots of individual EnR-CIPs analyzed in METABRIC and kmPlotter databases. Relapse-free survival of the 7 EnR-CIPs enriched in MCF-7-HER2 using kmPlotter and METABRIC datasets of luminal breast cancer patients who received ET.

**Figure S9.**
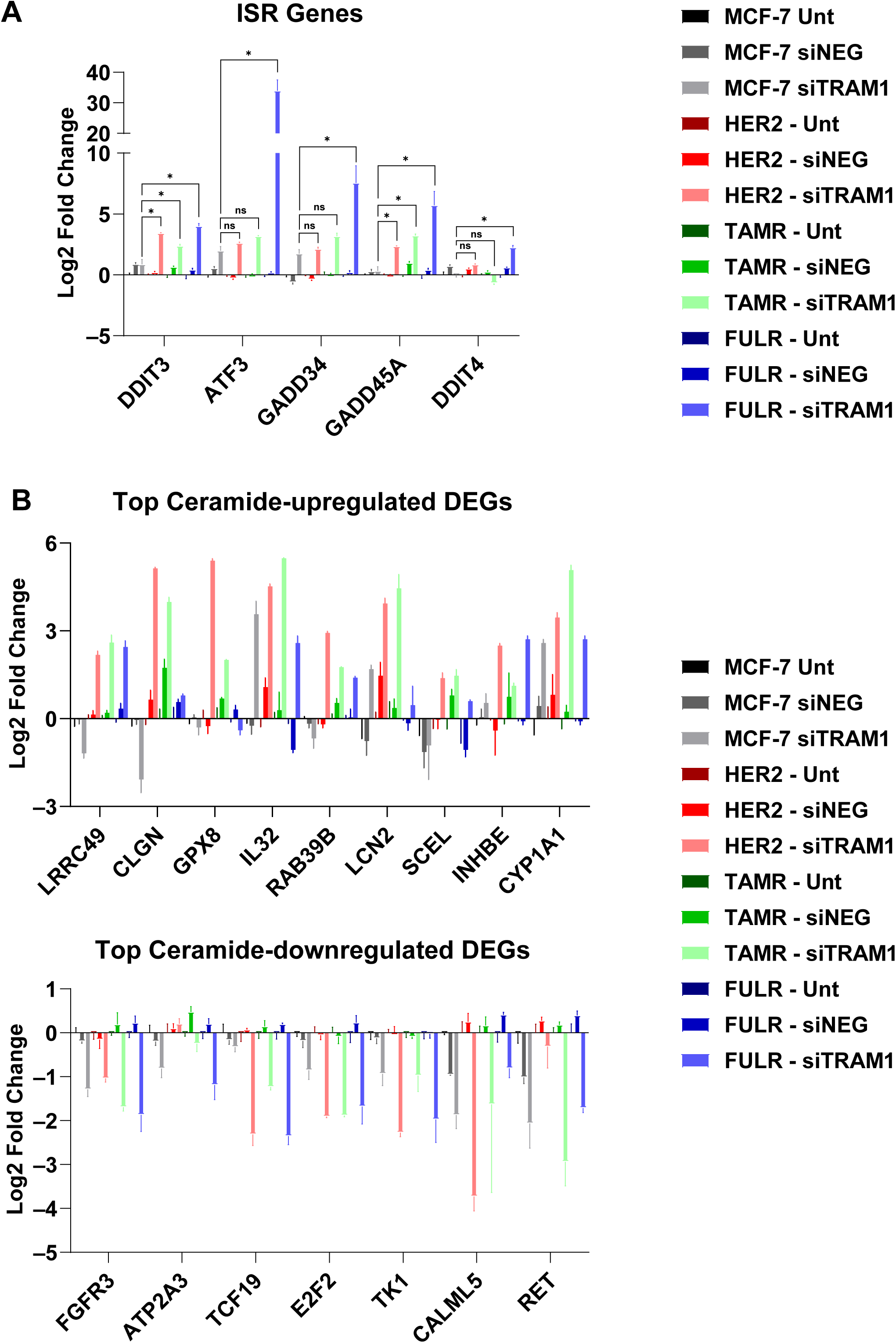
TRAM1 knockdown in ET-sensitive and -resistant MCF-7 cells. **A.** RT-qPCR of ISR genes after TRAM1 knockdown in ET-sensitive and ET-resistant MCF-7 cells. Data are presented as mean *±* SEM for triplicate samples from one experiment representative of three independent experiments. *P<0.05 by one-way ANOVA. **A.** RT-qPCR results showing expression of top C8-ceramide-induced (upregulated and downregulated) DEGs after TRAM1 knockdown. All Log_2_ fold changes are calculated relative to the untransfected controls. Data are presented as mean *±* SEM for triplicate samples from one experiment.

**Figure S10.**
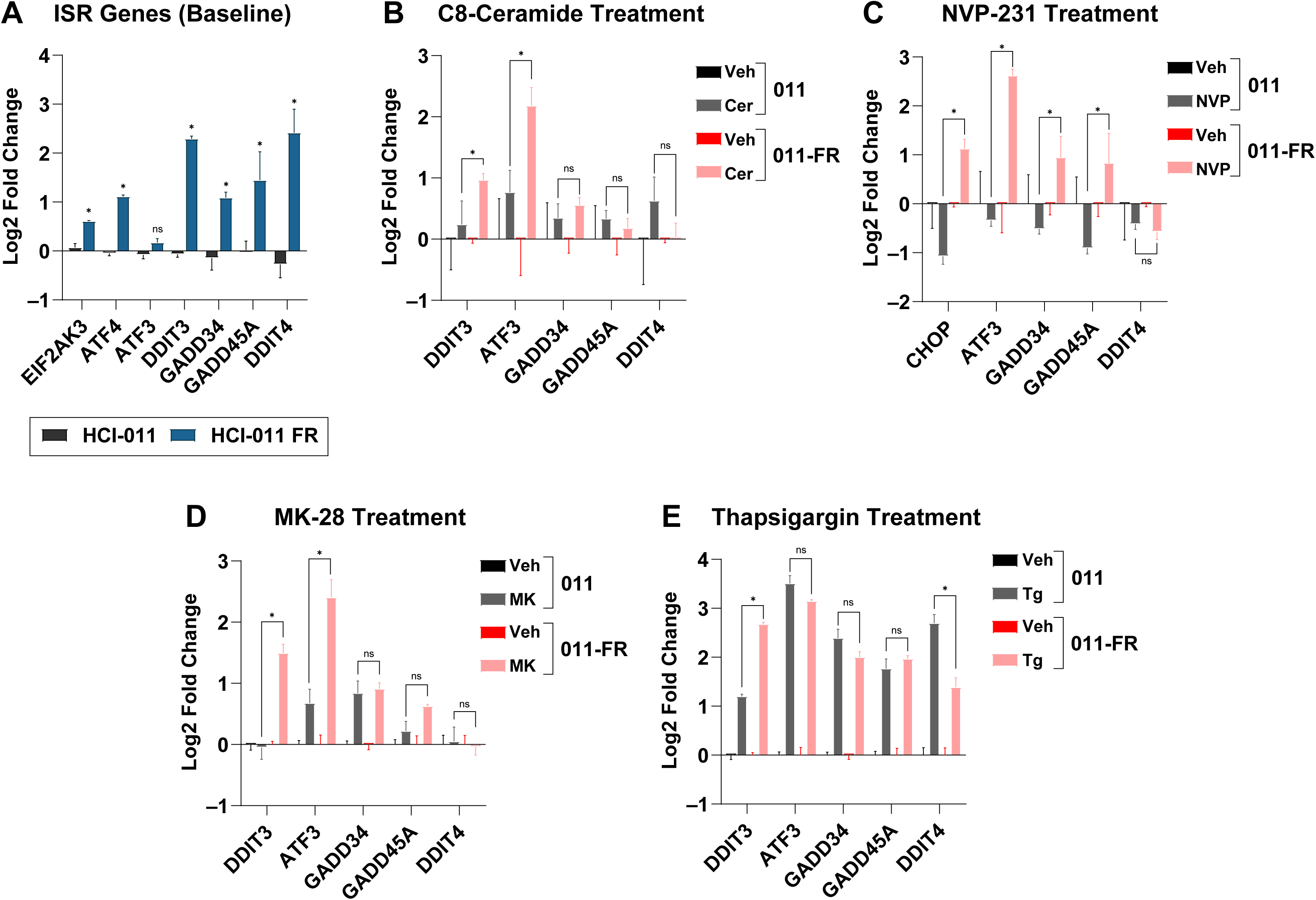
RT-qPCR experiments in PDxO models. **A.** RT-qPCR on ET-sensitive HCI-011 and its fulvestrant-resistant counterpart HCI-011-FR shows baseline expression of ISR genes. Data are presented as mean *±* SEM of triplicate samples from one experiment. All Log_2_ fold changes are calculated relative to parental HCI-011. *P<0.05 by unpaired Student’s t-test with Welch’s correction. **B-E.** Gene expression changes following 24-hour treatments with 10 µM C8-ceramide (**B**), 10 µM NVP-231 (**C**), 10 µM MK-28 (**D**), or 1 µM Thapsigargin (**E**). Data are presented as mean *±* SEM of triplicate samples from one experiment. *P<0.05 by unpaired Student’s t-test with Welch’s correction.

### TABLES

**Table S1** Differentially expressed genes induced by C8-Ceramide in each ET-sensitive and ET-resistant MCF-7 cell lines.

**Table S2** Pathway analysis of commonly altered genes by ceramides in all ET-resistant cells, performed using DAVID.

**Table S3** Ceramide-induced pathway changes in each ET-sensitive and ET-resistant MCF-7 cell line identified using GSEA.

**Table S4** Baseline alterations in cellular pathways across all ET-resistant models, as analyzed by GSEA.

**Table S5** Genes associated with Integrated Stress Response (ISR) pathway activation in multiple ET-resistant MCF-7 cell lines.

**Table S6** Significantly differentially abundant CIPs in MCF-7-HER2 cells compared to parental MCF-7 cells.

**Table S7** Ingenuity Pathway Analysis (IPA) showing the top upregulated and downregulated pathways in MCF-7-HER2 cells, based on differential expression of CIPs.

**Table S8** List of qRT-PCR primers used in this study.

## ACKNOWLEDGEMENT

Authors acknowledge Isra Alam, Daniel Lu (SUNY), Aanshi Vashi, Shreenithi Palamuthy, Kathryn Schart (UIC) for their help in performing experiments. Authors also thank UIC Genomic Research Core for RNA-seq and Dr. Seock-Won Youn (Viral Vector Core) for helping with cloning.

## AUTHOR CONTRIBUTION

Conceptualization: P.P., G.E.A.-G., J.F., and J.C.; Methodology: S.C., G.E.A.-G., and J.Q.; Validation: P.P. and J.C.; Formal analysis: P.P., S.C., G.K.S., F.A., H.Z., M.A., G.E.A.-G., and J.C.; Resources: R.S., D.T., and G.L.G.; Data Curation: P.P., S.C., H.Z., G.K.S., and F.A.; Writing — original draft preparation: P.P.; Writing — review and editing: P.P., S.C., G.K.S., F.A., D.T., R.S., G.L.G., G.E.A.-G. and J.C.; Supervision: G.E.A.-G. and J. C.; Project Administration, G.E.A.-G., J.F., J.C., and P.P.; Funding Acquisition, G.E.A.-G., J.F., J.C., and P.P.; All authors have read and agreed to the published version of the manuscript.

## INSTITUTIONAL REVIEW BOARD STATEMENT

All animal procedures were approved by the Office of Animal Care and Institutional Biosafety (OACIB) of University of Illinois Chicago, under protocol number 24-031, and followed relevant guidelines for the care and use of laboratory animals.

## INFORMED CONSENT STATEMENT

Not Applicable.

